# Role of Thalamus in Human Conscious Perception Revealed by Low-Intensity Focused Ultrasound Neuromodulation

**DOI:** 10.1101/2024.10.07.617034

**Authors:** Hyunwoo Jang, Panagiotis Fotiadis, George A. Mashour, Anthony G. Hudetz, Zirui Huang

## Abstract

The neural basis of consciousness remains incompletely understood. While cortical mechanisms of conscious perception have been extensively investigated in humans, the role of subcortical structures, including the thalamus, remains less explored. Here, we elucidate the causal contributions of different thalamic regions to conscious perception using transcranial low-intensity focused ultrasound (LIFU) neuromodulation. We hypothesize that modulating distinct thalamic regions alters perceptual outcomes derived from Signal Detection Theory. We apply LIFU to healthy human anterior (transmodal-dominant) and posterior (unimodal-dominant) thalamic regions, further subdivided into ventral and dorsal regions, during a near-threshold visual perception task. We show that high duty cycle modulation of the ventral anterior (VA) part of thalamus enhances object recognition sensitivity. Sensitivity enhancement magnitude correlates with the core-matrix cell compositions of the stimulated thalamic region. Connectivity analysis of a large-scale functional magnetic resonance imaging dataset confirms strong transmodal connectivity of VA thalamus with frontoparietal and default-mode networks. We also demonstrate target-invariant effects of high duty cycle LIFU disrupting object categorization accuracy. These findings provide causal insight into the cytoarchitectural and functional organization of the thalamus that shapes human visual experience, especially the role of matrix cell-rich, transmodal-dominant VA thalamus.

## INTRODUCTION

The neural basis of consciousness remains an active area of neuroscientific investigation^1–7^. While cortical mechanisms of conscious perception contents have been extensively studied^8–12^, the role of subcortical structures in encoding, enabling, or modulating such contents remains less explored^7,8,13^. With its intricate connections to the cortex, the thalamus is a central structure that is well-positioned to orchestrate conscious experience^13,14^. Traditionally, thalamic nuclei have been categorized into specific and nonspecific types^15^. In general, specific nuclei relay modal sensory information to designated cortical areas, whereas nonspecific nuclei project more diffusely and support cortical arousal or higher-order cognition^14,16^. For instance, the lateral geniculate nucleus, considered a specific nucleus, relays retinal input to the primary visual cortex^15^, while the nonspecific intralaminar thalamus is involved in controlling arousal, attention, and global states of consciousness^17–19^.

Recent studies suggest that cortical areas are functionally organized along a unimodal-to-transmodal continuum rather than distinct categories^20,21^. Within the cortex, unimodal areas process sensorimotor information, whereas transmodal areas integrate information and support complex cognitive functions. Building on this functional organization of the cortex, our previous work revealed a unimodal-transmodal gradient in thalamocortical connectivity^20^. This connectivity gradient features anterior and medial thalamic regions connecting predominantly with transmodal cortices, whereas posterior thalamic regions connect primarily with unimodal cortices. A similar gradient is also reflected by the cytoarchitecture of the thalamus: diffusely-projecting calbindin-expressing matrix cells are enriched anteriorly, and narrowly-projecting parvalbumin-expressing core cells are concentrated in posterior thalamus^20,22^. Notably, both the connectivity gradient and cytoarchitectural gradient support the hypothesis that the matrix cell-rich anterior thalamus is crucial for conscious perception^18,23–26^. Disruption of this region has also been linked to loss of consciousness^20^.

A promising approach to investigate the causal contribution of thalamic gradients to human consciousness, more specifically conscious sensory perception, is to selectively modulate the activity of specific thalamic subregions. Traditional methods such as deep-brain stimulation, optogenetics, and transcranial electric/magnetic stimulation are either invasive or lack sufficient spatial resolution^27,28^. In contrast, transcranial low-intensity focused ultrasound (LIFU; also referred to as tFUS) offers a non-invasive approach with higher precision for targeting deep structures^29–33^. Pioneering work has applied LIFU to human subcortical regions, including the thalamus, putamen, and basal ganglia^34–37^. Although the mechanisms underlying LIFU’s effects are still under investigation, with theories ranging from thermal effects^38^ to membrane pore formation^39^ and mechanosensitive channel activation^40^, the ability of LIFU to modulate neural activity and behavior is well-supported^29,39,41,42^. Nevertheless, LIFU’s potential for modulating human conscious perception and for probing its mechanisms, particularly in the visual domain, remains largely unexplored^43,44^.

LIFU has been shown to modulate neural activity in a bidirectional manner, with the directionality governed by sonication parameters such as the duty cycle (DC; the percentage of time the ultrasound is actively transmitted within each pulse cycle). Early work using neuroimaging, electrophysiological, and behavioral measures indicates that higher DC values generally elicit excitatory effects, whereas lower DC values result in inhibition^33–35,39,40,45,46^. This pattern is supported by both computational models and *in vivo* experiments, which demonstrate that 70% DC effectively lowers neuronal activation thresholds and facilitates excitation^40,47–49^. In contrast, 5% DC is associated with diminished evoked potentials and suppressed blood oxygen level-dependent (BOLD) responses consistent with functional inhibition^40,49–51^. These observations align with the neuronal intramembrane cavitation excitation model, which posits that the value of DC determines the direction of neuromodulation^39,47,49^.

In this work, we aim to elucidate the causal roles of human thalamus in conscious visual perception by applying LIFU with region- and parameter-specific precision. We hypothesize that targeting distinct thalamic regions—each positioned along the unimodal-to-transmodal gradient— and varying the sonication duty cycle will lead to distinct changes in perceptual outcomes. To test this hypothesis, we sequentially modulate four gradient-defined thalamic areas^52^ (ventral anterior, ventral posterior, dorsal anterior, and dorsal posterior) while healthy human volunteers perform a near-threshold visual task. Perceptual changes are quantified using Signal Detection Theory (SDT) metrics. To interpret the observed effects at cytoarchitectural and network levels, we conduct complementary analyses with core-matrix cell composition and thalamocortical connectivity data. Our findings reveal that high duty cycle modulation of the matrix cell-rich, transmodal-dominant ventral anterior thalamus selectively enhances object recognition sensitivity. Additionally, we demonstrate target-invariant and duty cycle-dependent effects on the accuracy of object categorization, observed regardless of the target region.

## RESULTS

Sixty participants (age: 25.9 ± 6.3, mean ± SD; 38 females, 22 males) were randomly assigned to one of two duty cycle (DC) groups. One group received LIFU with a high duty cycle (70%), while the other received LIFU with a low duty cycle (5%) with equal spatial-peak temporal-average intensity of *I_spta_* = 0.72 W cm^-2^ (Fig. 1a). Six participants were excluded from analysis due to technical issues, leaving 54 participants in the final analysis (27 in each group; Supplementary Fig. 1).

**Fig. 1:**
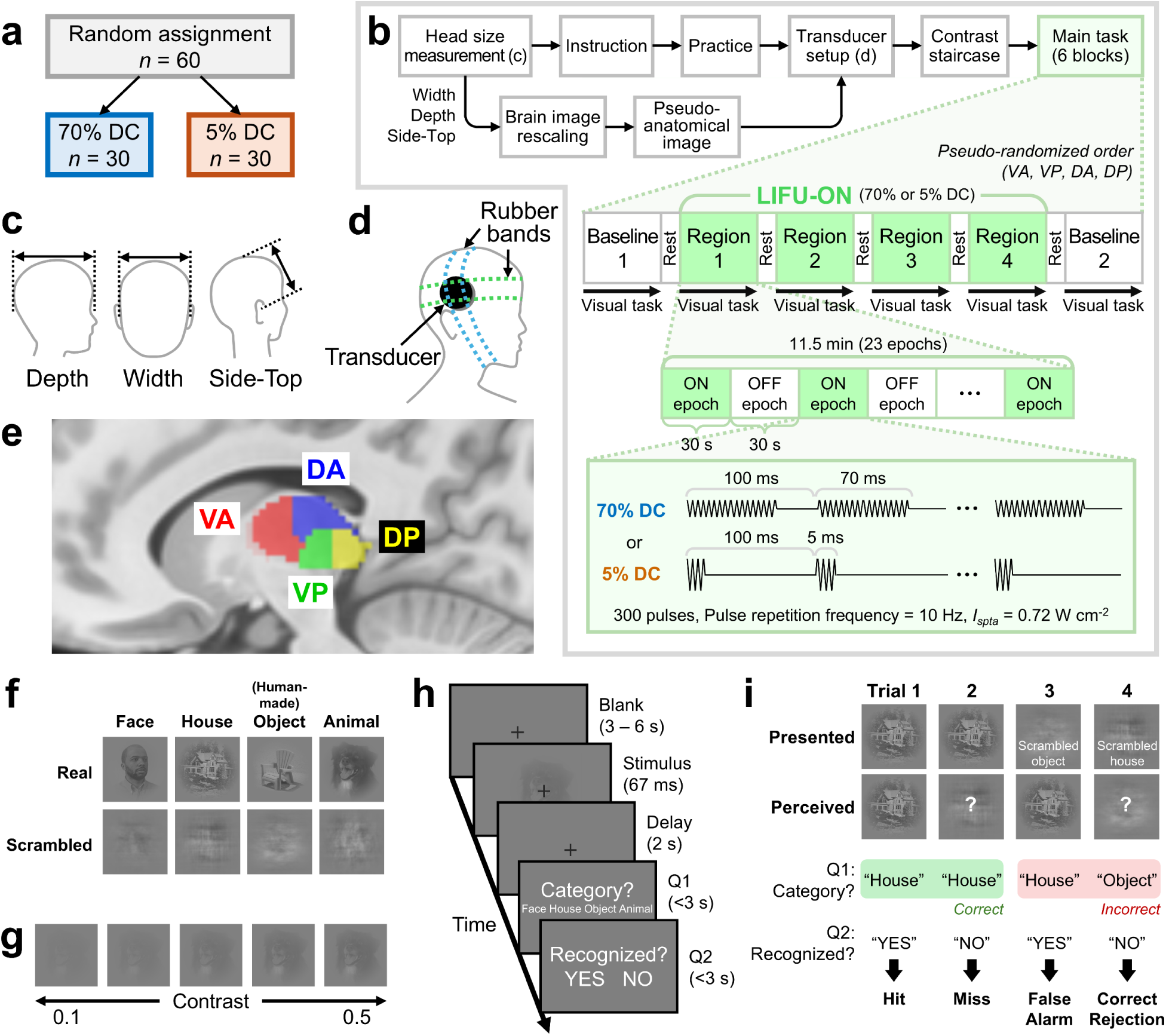
Overview of the block design, experimental setup, and behavioral paradigm. **a** Randomized study group assignment. Sixty subjects were randomly assigned to one of two groups, each consisting of 30 participants, corresponding to either a 70% or 5% DC. 54 participants (27 per group) were included in the final analysis. **b** The timeline of experimental sessions, including initial preparation steps and the main task. The main task included two outer baseline blocks and four inner LIFU-ON blocks with a pseudo-randomized order of targeting regions. Each block comprised 100 trials. Each LIFU-ON block included 23 alternating 30-second ON and OFF epochs (total 11.5 min). Visual tasks were conducted continuously within each block. Each ON epoch consists of 300 pulses (pulse repetition frequency = 10 Hz) at a fixed DC and temporal-average intensity (*I_spta.3_* = 0.72 W cm^-2^). The pulse-averaged intensity was higher at 5% DC. **c** Measurement locations for head size, including width, depth, and side-to-top. **d** Illustration of the transducer fixed by two perpendicular rubber bands on the participant’s head. **e** Sagittal view for the four target regions, including left VA, VP, DA, and DP. **f** Examples of real and scrambled images for each category (face, house, human-made object, and animal). **g** Example of an animal image displayed at different contrast levels. **h** Trial structure showing the sequence of events: a blank period, stimulus presentation, delay, and two questions about categorization and recognition. **i** Illustration of the four trial types (hit, miss, false alarm, and correct rejection). Due to copyright limitations, the actual images used in our experiment are not shown. Copyright-free images included in this figure were obtained from https://www.pexels.com. DC: duty cycle; VA: ventral anterior thalamus; VP: ventral posterior thalamus; DA: dorsal anterior thalamus; DP: dorsal posterior thalamus.

The experiment involved key steps including transducer setup, image contrast titration, and six blocks of the visual task (Fig. 1b-d). There were two outer LIFU-OFF baseline blocks (Baseline-1 and Baseline-2) and four inner LIFU-ON blocks. During each LIFU-ON block, LIFU was directed toward one of four regions of the left thalamus, including the ventral anterior (VA), ventral posterior (VP), dorsal anterior (DA), and dorsal posterior (DP) thalamus (Fig. 1e; see Supplementary Table 1 for sonication parameter summary; see Methods: Target locations for exact coordinates). The order of target regions was pseudo-randomized and counterbalanced among participants, and each participant received a consistent duty cycle throughout the session.

We employed a well-established near-threshold visual object recognition and categorization task^8,9^. Four object categories were used: faces, houses, human-made objects, and animals (Fig. 1f). Before the main task, using an adaptive staircase paradigm, we individually titrated the image contrast (Fig. 1g) to determine the threshold of conscious perception (i.e., a contrast yielding 50% recognition rate)^8^. Here, “recognition” was operationally defined as the perception of an object that makes sense in the real world, as opposed to meaningless noise-like patterns. During each trial, participants responded to two questions: one regarding the category of the presented image (i.e., “Which category does the image belong to?”) and another concerning their subjective perceptual experience (i.e., “Did you recognize the image?”) (Fig. 1h).

Importantly, we also included scrambled images, which comprised 20 out of 100 trials. Therefore, the recognition rates for real and scrambled images could be interpreted as hits and false alarms, respectively (Fig. 1i). This enabled us to measure sensitivity (*d’*) and decision bias (quantified by criterion *c*) of object recognition within the SDT framework (see Methods: Signal detection theory analysis). Sensitivity reflects the ability to discriminate real and scrambled images, and higher criterion signifies a tendency to say “NO” (i.e., “I didn’t recognize the image”) to the Question-2, reflecting a conservative stance. Categorization accuracy was also quantified for real and scrambled images, independently of object recognition.

Before LIFU was administered (i.e., at Baseline-1), participants recognized 53.3 ± 16.1% (mean ± SD) of the real images (Supplementary Fig. 2a), which was not significantly different from the intended 50% recognition rate, confirming a successful implementation of the staircase procedure. See SI text: Perceptual outcomes at Baseline-1 and Supplementary Fig. 2 for detailed analysis of baseline perceptual outcomes.

We examined whether LIFU induced any carry-over effects within our approximately one-hour experimental timeframe by comparing two outer LIFU-OFF blocks (Baseline-1 and Baseline-2). For both duty cycles, we found no significant difference in any perceptual outcomes (*p* > 0.05, Wilcoxon signed-rank test, two-sided; Supplementary Fig. 3). These results suggest that LIFU did not produce detectable long-lasting or cumulative effects on visual perception in our experimental condition.

### Ultrasound beam analysis and targeting accuracy

For each LIFU-ON block, we assessed various beam profiles, including targeting accuracy, beam shape, attenuation by tissue, and thermal effects (Fig. 2a). As proof of concept, we used a pseudo-anatomical reference for beam targeting and post-hoc analysis (see Methods: Template rescaling method for details). This was achieved by rescaling the MNI template image to match each participant’s head size measured before the main task (Fig. 1c). We validated this approach on a separate dataset (*n* = 13) by comparing actual anatomical images with their rescaled pseudo- anatomical images, finding an average deviation of 2.15 ± 0.82 mm (mean ± SD; Supplementary Fig. 4).

**Fig. 2:**
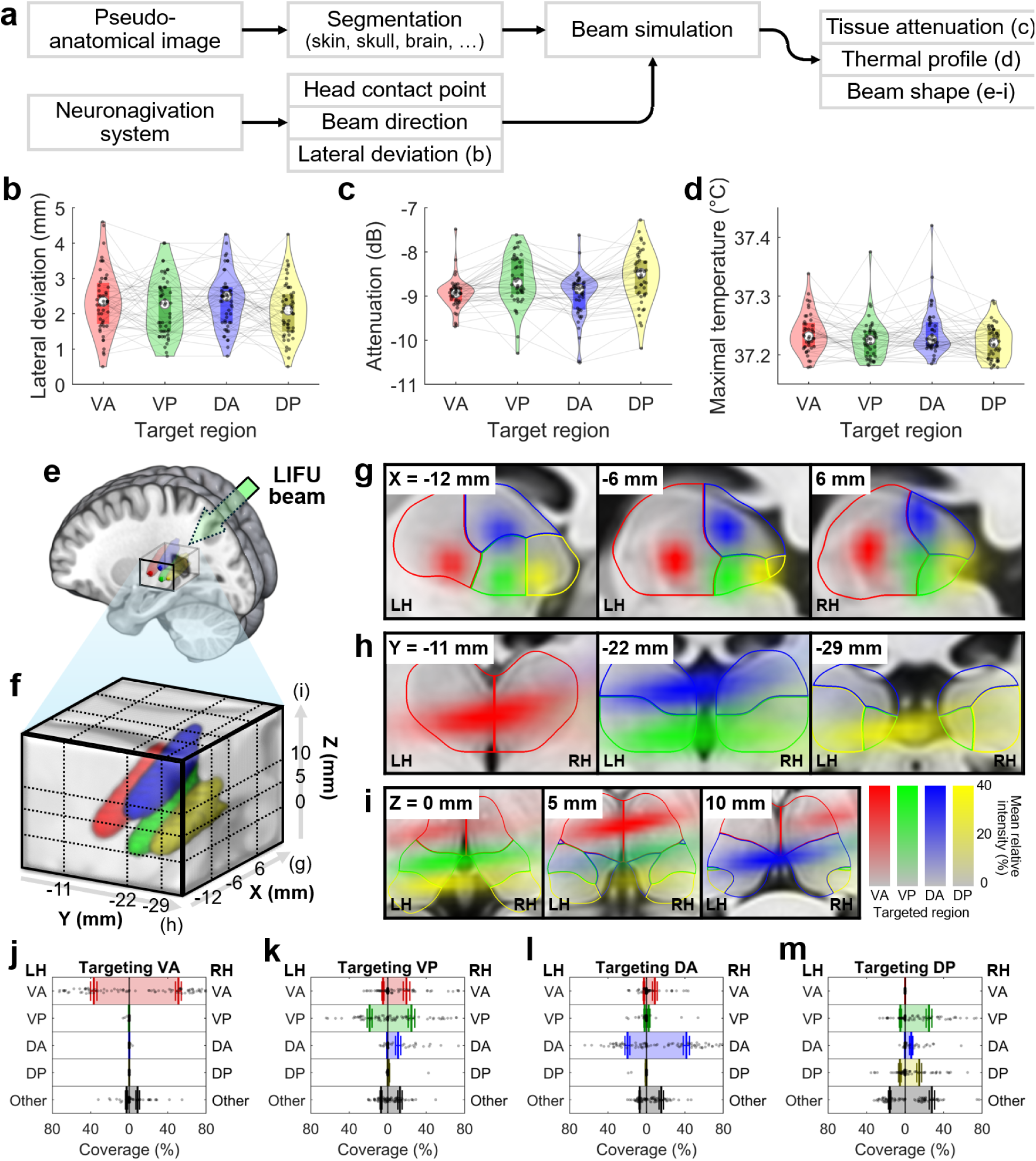
Post-hoc LIFU beam analysis and targeting accuracy. **a** Flowchart summarizing the beam analysis and simulation pipeline. **b** Lateral deviation of the beam from its target, defined as the Euclidean distance between the beam vector and the target coordinate. **c, d** Simulated **c** tissue-induced attenuation and **d** maximal tissue temperature. In panels b-d, boxes indicate interquartile ranges, and white circles show median values. **e** Three-dimensional rendering of the mean relative beam intensity across participants. Colored regions highlight voxels where the mean relative intensity exceeds 10%. An arrow indicates the LIFU beam originating from the right side. **f** Zoomed-in view of panel e, with dashed lines indicating the locations of cross-sections. **g-i** Cross-sectional views in the **g** sagittal, **h** coronal, and **i** axial planes, displaying mean relative intensity and outlines of bilateral thalamic regions. Higher opacity reflects greater intensity. **j-m** Bilateral coverage percentages for beams targeting left **j** VA, **k** VP, **l** DA, and **m** DP. The coverage percentage indicates how much of a “beam-present” volume (defined by > 50% intensity) falls within the thalamic (or other) regions. Boxes indicate mean values, and error bars show standard error. For panel b, VA: *n* = 51; VP: *n* = 50; DA: *n* = 53; DP: *n* = 54. For panels c-m, VA: *n* = 44; VP: *n* = 44; DA: *n* = 45; DP: *n* = 46. Color coding: VA (red), VP (green), DA (blue), and DP (yellow). VA: ventral anterior thalamus; VP: ventral posterior thalamus; DA: dorsal anterior thalamus; DP: dorsal posterior thalamus; LH: left hemisphere; RH: right hemisphere. Source data are provided as a Source Data file.

In a free-field condition, our transducer produces a bullet-like beam shape with an aspect ratio ≈ 9, measuring 30.3 mm in length and 3.3 mm in width (defined by -3 dB attenuation; Supplementary Fig. 5). Given this elongated shape, the lateral deviation reported by the neuronavigation system is a straightforward measure of the beam orientation accuracy. Here, lateral deviation is defined as the Euclidean distance between the beam vector (originating from the transducer center) and the target coordinate. The mean lateral deviation was 2.25 ± 0.81 mm (mean ± SD; Fig. 2b) and did not differ significantly across regions (*p* = 0.2155, Friedman’s test; see Source Data for statistics), with maximal deviation not exceeding 5 mm, together confirming that the beam was successfully oriented toward the targets.

We then simulated the beam on the segmented pseudo-anatomical image *in silico*. In line with the literature^53–55^, we observed substantial intensity attenuation of -8.78 ± 0.56 dB (mean ± SD; Fig. 2c), equating to approximately 87% energy loss. This yielded estimated pressure of 72.6 ± 4.2 kPa for 70% DC and 258.2 ± 18.3 kPa for 5% DC (mean ± SD; Supplementary Fig. 6). This pronounced energy loss is mainly attributable to skull^53,54,56,57^. The maximal temperature rise simulated across skin, skull, and brain was 0.23 ± 0.04 °C (mean ± SD) and didn’t exceed 0.5 °C (Fig. 2d), complying with ITRUSST risk recommendations^58^. Additionally, the beam depth (i.e., the Euclidean distance from the head-transducer contact point to the location of maximal simulated intensity) was simulated as 67.4 ± 3.4 mm (mean ± SD; Supplementary Fig. 7), approximately 15% shorter than the free-field focal depth of 80.7 mm (Supplementary Table 1,2), consistent with skull-induced beam shortening reported previously^55^.

Next, we evaluated how LIFU beams covered each thalamic region, as visualized by the mean relative intensity across subjects (Fig. 2e-i; see Supplementary Fig. 8-10 for individual trajectories). Because the LIFU beam originated from the right temple (Fig. 2e) and was effectively shortened by the skull (Supplementary Fig. 7), we observed significant bilateral coverage for all four target regions (Fig. 2g-i). To quantify this further, we computed a coverage ratio defined by how much of the “beam-present” volume (defined by > 50% intensity) belongs to the thalamic (or other non-thalamic) regions (Fig. 2j-m). We found excellent beam localization in bilateral VA, VP, and DA (Fig. 2j-l). In the case of DP targeting, the beam partially encompassed bilateral VP, presumably because the DP regions lie lateral to VP (Fig. 2m). Overall, these post-hoc analyses confirm successful targeting of each thalamic region.

### Effects of LIFU administration on perceptual outcomes

To determine whether LIFU-targeting of different thalamic regions produces differential effects on visual perception, we conducted omnibus *F*-tests using linear mixed-effects (LME) models for each perceptual outcome and duty cycle condition. Following the observation that the two baseline conditions did not differ significantly (Supplementary Fig. 3), we averaged them into a single baseline measure, yielding five targeting conditions (baseline plus four thalamic targets) for comparison.

The omnibus analysis revealed significant regional effects for three conditions (Table 1). At 70% DC, we observed significant omnibus effects for sensitivity *d’* (*F*(4,90) = 4.6129, *p*_FDR_ = 0.0118) and real-image accuracy (*F*(4,107) = 5.4470, *p*_FDR_ = 0.0060). At 5% DC, criterion *c* showed a significant omnibus effect (*F*(4,77) = 4.1100, *p*_FDR_ = 0.0181). Other perceptual measures, including hit rate, false alarm rate, and scrambled image accuracy, did not demonstrate significant effects after FDR correction at either duty cycle condition. To ensure that these effects were not artifacts of baseline averaging, we conducted control analyses using each baseline condition separately, showing that the main findings remained robust across both individual baselines (Supplementary Table 3). We also confirmed that none of the nuisance variables included in our LME model showed significant effects in omnibus ANOVA (all *p*_FDR_ > 0.05; Supplementary Table 4; see Methods: Statistics for details).

**Table 1:** Omnibus ANOVA for the effect of sonication target region on behavioral outcomes.

| DC | Perceptual outcome | df<sub>1</sub> | df<sub>2</sub> | F-value | p-value | FDR-corrected p-value |
| --- | --- | --- | --- | --- | --- | --- |
| 70% | Hit rate | 4 | 107 | 1.3407 | 0.2596 | 0.5192 |
|  | FA rate | 4 | 110 | 0.6525 | 0.6263 | 0.8351 |
| | Sensitivity $d'$ | 4 | 90 | 4.6129 | 0.0020 | **0.0118** |
| | Criterion $c$ | 4 | 89 | 0.5325 | 0.7122 | 0.8546 |
|  | Real image accuracy | 4 | 107 | 5.4470 | 0.0005 | **0.0060** |
|  | Scrambled image accuracy | 4 | 111 | 0.2665 | 0.8989 | 0.9553 |
| 5% | Hit rate | 4 | 104 | 2.3734 | 0.0570 | 0.1710 |
|  | FA rate | 4 | 104 | 2.0005 | 0.0999 | 0.2398 |
| | Sensitivity $d'$ | 4 | 78 | 0.7452 | 0.5641 | 0.8351 |
| | Criterion $c$ | 4 | 77 | 4.1100 | 0.0045 | **0.0181** |
|  | Real image accuracy | 4 | 103 | 0.9165 | 0.4574 | 0.7841 |
|  | Scrambled image accuracy | 4 | 106 | 0.1658 | 0.9553 | 0.9553 |

Significant values (FDR-corrected $p < 0.05$ ) are highlighted as red. DC: duty cycle; df: degree of freedom; FA: false alarm.

For the three conditions that demonstrated significant omnibus effects, we conducted post-hoc pairwise comparisons to identify which specific thalamic targets potentially drove the observed regional differences (Fig. 3; see Supplementary Fig. 11 for all other perceptual outcomes).

**Fig. 3:**
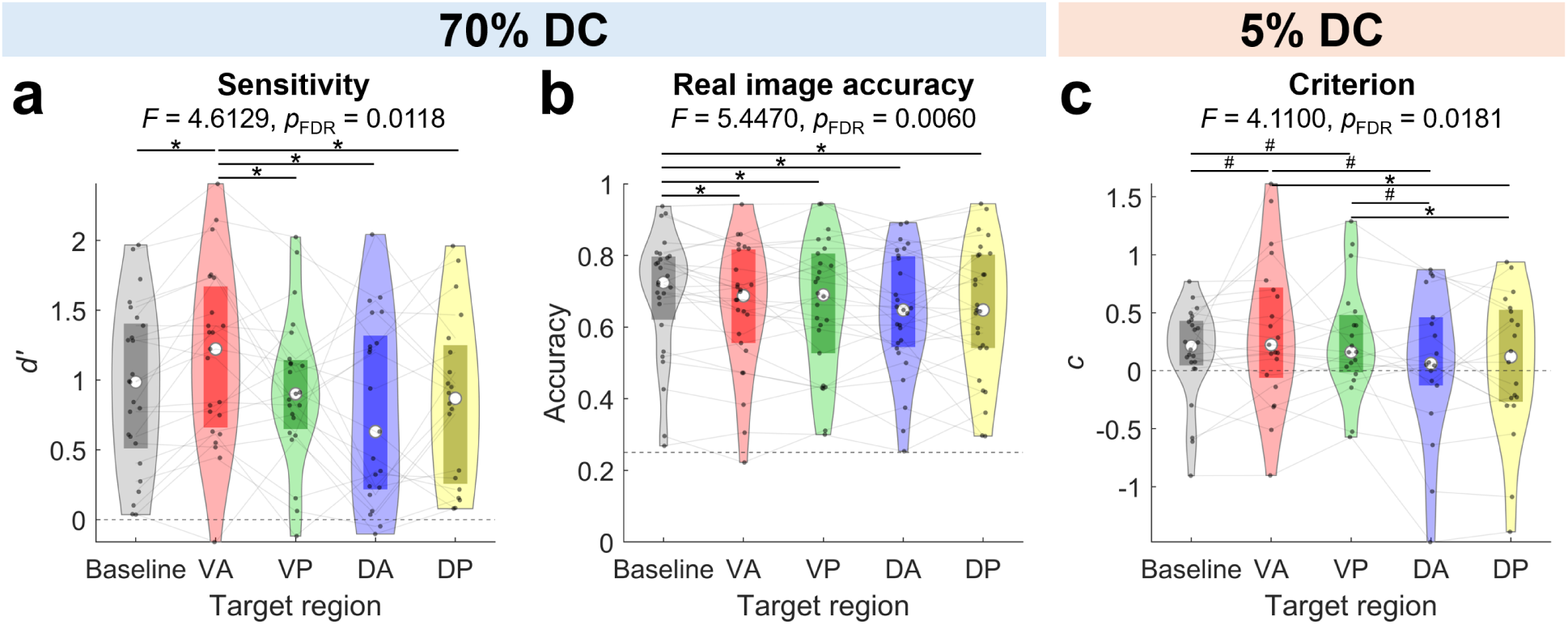
Target-specific changes in perceptual outcomes during human thalamic LIFU application. Perceptual outcomes that demonstrated significant omnibus effects, measured at **a-b** 70% and **c** 5% DC, including **a** sensitivity *d’*, **b** categorization accuracy for real images, and **c** criterion *c*, are shown. Asterisks and pounds denote significant differences in post-hoc pairwise comparisons (asterisk: FDR-corrected *p* < 0.05; pound: FDR-uncorrected *p* < 0.05). Boxes indicate interquartile ranges. Median values are marked by white circles. Dashed lines are reference values for each measure: no discrimination of *d’* = 0 (a), chance accuracy of 0.25 (b), and no bias of *c* = 0 (c). For panel a, Baseline: *n* = 25; VA: *n* = 23; VP: *n* = 23; DA: *n* = 21; DP: *n* = 20; for panel b, Baseline: *n* = 27; VA: *n* = 26; VP: *n* = 26; DA: *n* = 27; DP: *n* = 27; for panel c, Baseline: *n* = 24; VA: *n* = 21; VP: *n* = 19; DA: *n* = 18; DP: *n* = 20. DC: duty cycle; FA: false alarm; VA: ventral anterior thalamus; VP: ventral posterior thalamus; DA: dorsal anterior thalamus; DP: dorsal posterior thalamus. Statistics and source data are provided as a Source Data file.

At 70% DC, LIFU targeting of the VA nucleus specifically enhanced sensitivity *d’* compared to baseline (VA vs. baseline: *p*_FDR_ = 0.0246; paired Cohen’s *d* = 0.9164) and significantly outperformed all other target regions (Fig. 3a; VA vs. VP: *p*_FDR_ = 0.0246, VA vs. DA: *p*_FDR_ = 0.0034, VA vs. DP: *p*_FDR_ = 0.0119; see Supplementary Table 5 for full statistics). This pattern demonstrates that the omnibus effect was driven primarily by VA-specific enhancement rather than a generalized effect.

We also identified that the increase in sensitivity during VA thalamus at 70% DC sonication became more robust as the lateral deviation of the beam decreased (Supplementary Fig. 12). Specifically, the height of the fitted Gaussian curve was significantly positive for 70% DC (*p*_FDR_ = 0.0286) but not for 5% DC (*p*_FDR_ = 0.7679; see Source Data for full statistics). These findings support the target-specificity of this effect and validate our targeting precision.

For categorization accuracy of real images, a consistent decrease across all four thalamic regions relative to baseline was observed (Fig. 3b; baseline vs. all four regions: *p*_FDR_ < 0.05; average paired Cohen’s *d* = -0.4545; Supplementary Table 5), suggesting a target-invariant disruptive effect of high-DC thalamic sonication on visual categorization tasks.

At 5% DC, the effects of LIFU on criterion *c* revealed a more complex pattern. We observed elevated criterion (i.e., shift toward a conservative stance) in both VA and VP sonication compared to baseline and to dorsal targets (DA and DP), although many of these pairwise comparisons reached only marginal significance after FDR correction (Fig. 3c; VA vs. DP and VP vs. DP: *p*_FDR_ < 0.05; Supplementary Table 5). This nuanced pattern suggests regionally heterogeneous effects that are not uniformly robust across all comparisons.

### Cytoarchitectural correlates of LIFU-induced perceptual modulation

To test whether thalamic cytoarchitecture predicts the observed perceptual effects, we conducted correlation analyses between the core-matrix cell composition of stimulated regions and behavioral outcomes. Core and matrix cells can be distinguished by their expression of calcium-binding proteins, with core cells expressing higher levels of Parvalbumin (PVALB) and matrix cells expressing higher levels of Calbindin (CALB1)^22^. We quantified the relative composition of these cell types using the CP_T_ metric, as defined by the difference between CALB1 and PVALB expression levels derived from the Allen Human Brain Atlas, which can be used as a proxy for relative matrix cell richness^20,22,59^, i.e., a higher CP_T_ value indicates a more matrix-cell rich area.

For each LIFU-ON block, we calculated the mean CP_T_ value across beam-present voxels (> 50% relative intensity; black contours in Supplementary Fig. 8-10). We then correlated these CP_T_ values with baseline-subtracted changes in the three perceptual outcomes that survived omnibus testing. The analysis revealed a significant positive correlation between CP_T_ and change in sensitivity *d’* at 70% DC (Spearman’s *ρ* = 0.3479, *p*_FDR_ = 0.0135; Fig. 4a), indicating that regions with higher matrix cell density produced greater enhancement in perceptual sensitivity when sonicated by high DC. In contrast, no significant correlations emerged for real image accuracy at 70% duty cycle (*ρ* = -0.1026, *p*_FDR_ = 0.3590; Fig. 4b) or criterion *c* at 5% duty cycle (*ρ* = 0.1748, *p*_FDR_ = 0.2304; Fig. 4c, see Source Data for full statistics).

**Fig. 4:**
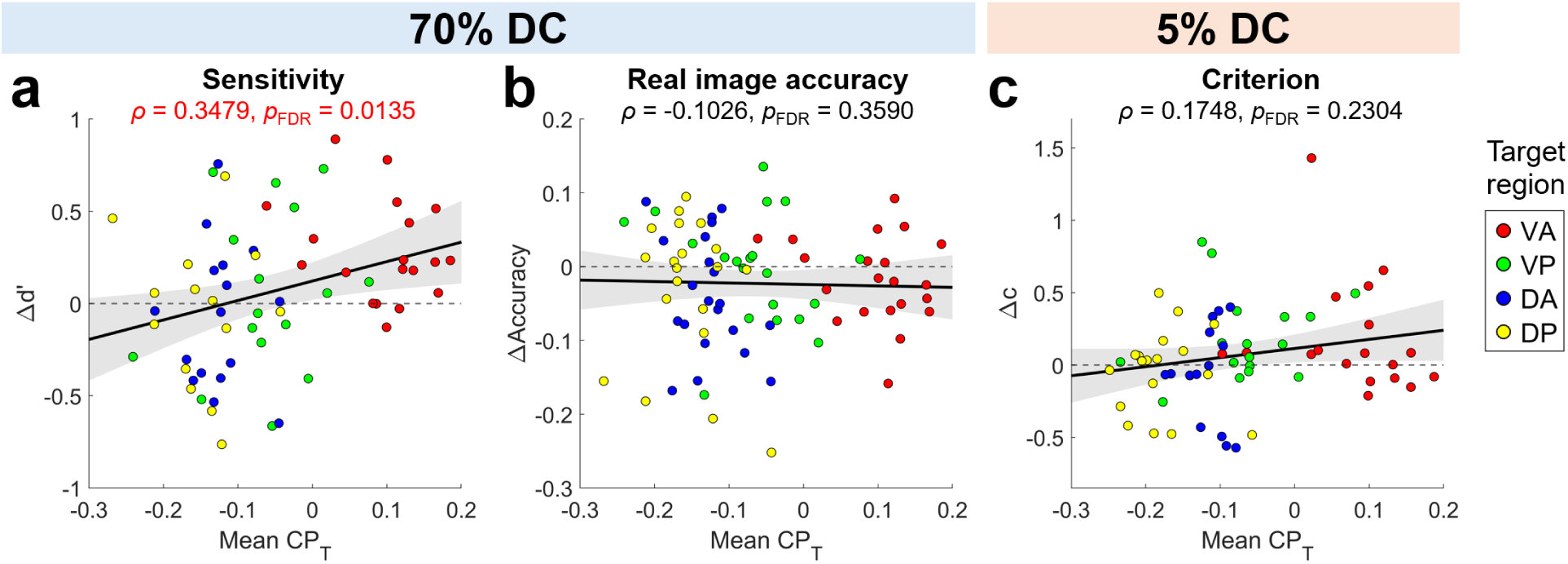
Correlation between matrix cell richness at beam-present regions (*x*-axis) and behavioral outcome changes (*y*-axis) across LIFU-ON blocks. CP_T_ reflects the relative difference between Calbindin (CALB1) and Parvalbumin (PVALB) expressions. Positive and negative CP_T_ values indicate matrix cell-richness and core cell-richness at the beam center (i.e., voxels exceeding 50% relative intensity), respectively. Spearman’s *ρ* and FDR-corrected *p*-values are provided. Circle colors indicate target regions. Solid lines and shades show linear fits and 95% confidence intervals. VA: ventral anterior thalamus; VP: ventral posterior thalamus; DA: dorsal anterior thalamus; DP: dorsal posterior thalamus. Statistics and source data are provided as a Source Data file.

### Functional connectivity between thalamic regions and the cortex

To gain deeper insights into the network-level neural correlates of both target-specific and target-invariant effects of thalamic LIFU, we analyzed the connectivity profiles of the four thalamic regions by utilizing a large-scale functional magnetic resonance imaging (fMRI) dataset of the Human Connectome Project (*n* = 1009)^60^.

First, we assessed the connectivity profiles of the four thalamic regions at the network level (Fig. 5a,b). For each thalamic region of interest (ROI), we calculated the functional connectivity with 400 cortical ROIs and averaged their percentile rank across seven canonical networks^61^. In line with our prior research^20^, we observed a graded shift in connectivity profiles along a unimodal-transmodal gradient. The VA thalamus displayed transmodal dominance, exhibiting higher connectivity with frontoparietal and default-mode networks (red curve in Fig. 5b) compared to the other three thalamic regions. This was followed by DA thalamus, with a trend toward unimodal dominance observed in DP and VP thalamus.

**Fig. 5:**
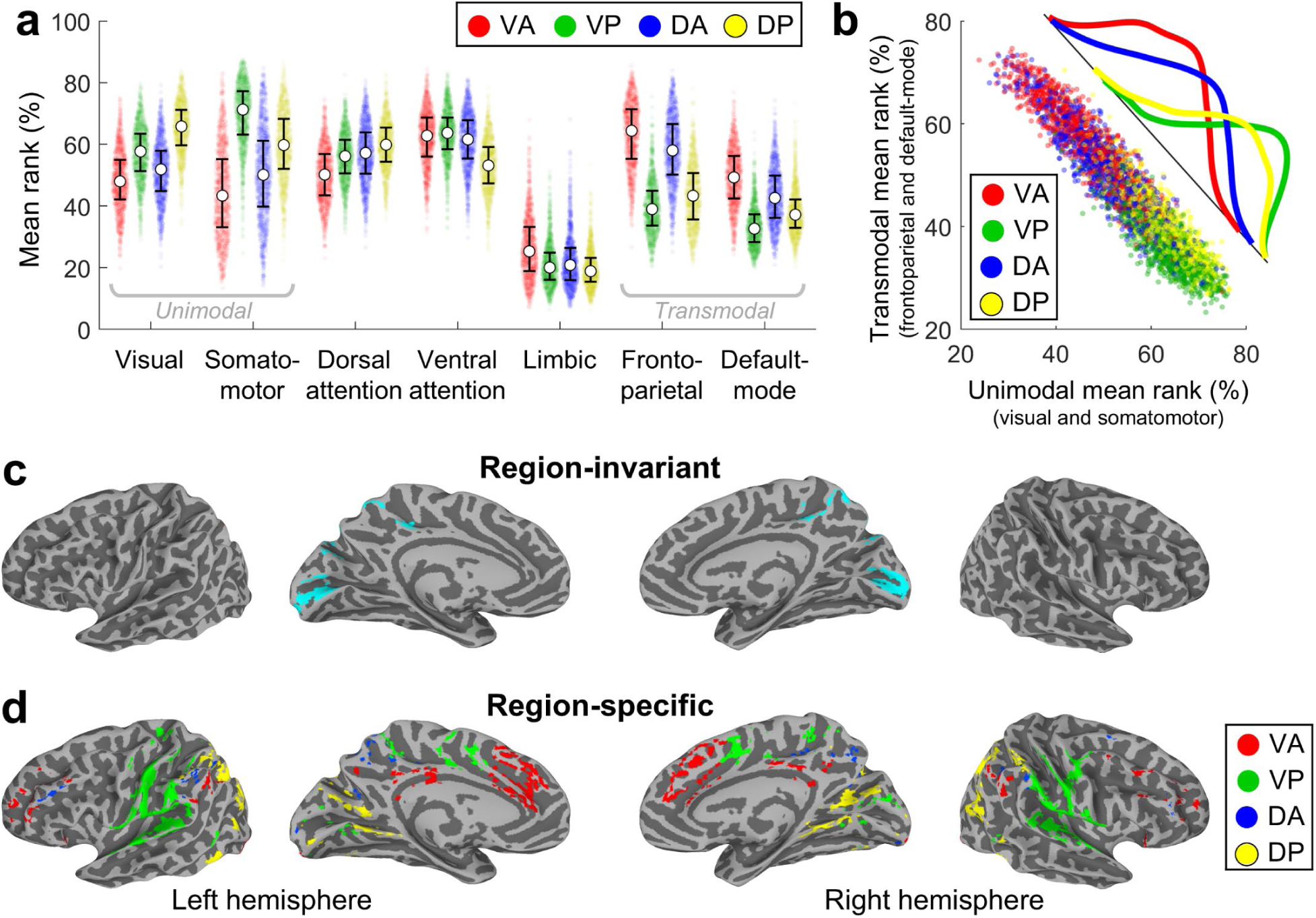
Functional connectivity profiles between four thalamic regions and the cortex. **a, b** Network-level analysis. **a** Percentile ranks of functional connectivity of 400 cortical regions and each thalamic region, averaged within seven canonical Schaefer-Yeo networks (visual, somatomotor, dorsal attention, ventral attention, limbic, frontoparietal, and default-mode). Each dot represents an individual (*n* = 1009). White circles indicate median values and error bars denote interquartile ranges. **b** Distribution of mean connectivity ranks of unimodal (visual and somatomotor) vs. transmodal (frontoparietal and default-mode) networks with each thalamic region. Curves represent kernel-smoothed histograms across the direction of maximal variance. **c, d** Voxel-based analysis. Cortical voxels exhibiting the top 10% strongest connectivity with each thalamic region as a seed, that are **c** common across four thalamic regions (i.e., region-invariant) and **d** unique to each region (i.e., region-specific). VA: ventral anterior thalamus; VP: ventral posterior thalamus; DA: dorsal anterior thalamus; DP: dorsal posterior thalamus. Source data are provided as a Source Data file.

Second, we examined the thalamocortical connectivity at a finer spatial scale, specifically at voxel level. We retained the cortical voxels exhibiting the top 10% strongest connectivity with each respective thalamic region as a seed. The connectivity profiles of the four regions showed considerable overlap, particularly within the more primary visual cortex (cyan regions in Fig. 5c). Region-specific connectivity patterns were also observed. The VA thalamus was predominantly connected with the medial prefrontal cortex (mPFC) and dorsolateral prefrontal cortex (dlPFC) (red regions in Fig. 5d). The VP thalamus exhibited unique connectivity with the somatomotor and auditory cortices (green regions in Fig. 5d, see also high connectivity with somatomotor network in Fig. 5a). The DP thalamus showed unique clusters within various regions in the visual cortex (yellow regions in Fig. 5d). The connectivity profile of the DA thalamus largely overlapped with the VA thalamus (Supplementary Fig. 13), with only a few unique clusters located in left dlPFC and right temporal-parietal junction (blue regions in Fig. 5d).

## DISCUSSION

This study aimed to causally probe how different thalamic regions contribute to conscious perception in humans. Through low-intensity focused ultrasound (LIFU) neuromodulation, we identified a specific role for the matrix cell-rich, transmodal-dominant ventral anterior (VA) thalamus in modulating the sensitivity of conscious visual perception (Fig. 6a). Sonicating the VA thalamus with a high duty cycle (DC) significantly increased object recognition sensitivity, consistent with the strong functional connectivity of the matrix cell-rich VA thalamus with transmodal cortical areas, particularly the medial and dorsolateral prefrontal cortices. Additionally, we observed target-invariant decrease in object categorization accuracy under high DC sonication (Fig. 6b). Together, these findings demonstrate the causal involvement of matrix cell-rich thalamic regions in conscious perception and underscore the modulatory potential of thalamocortical networks in shaping visual experience.

**Fig. 6:**
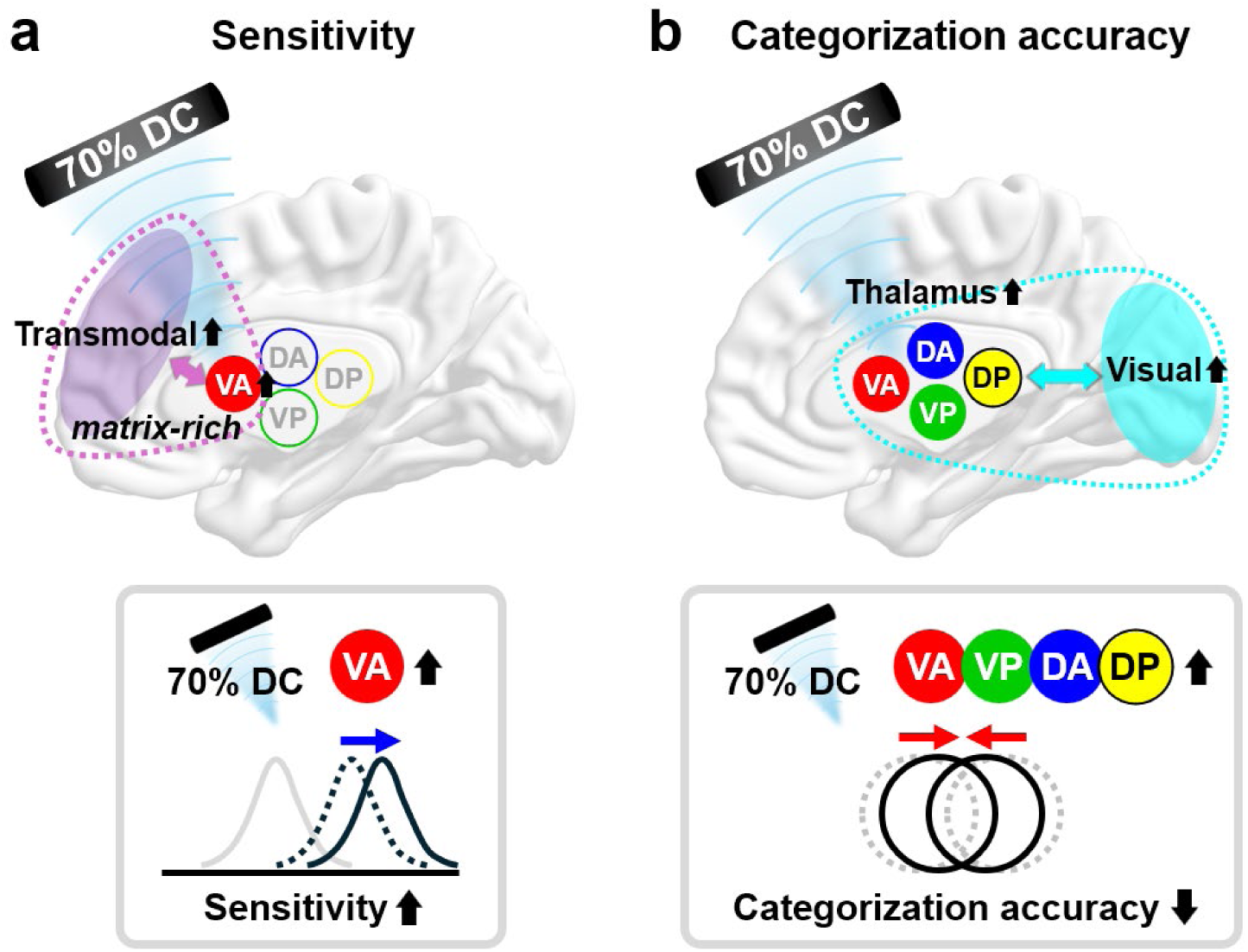
Summary of thalamic LIFU effects on human visual perception. **a** Sensitivity increases when the VA thalamus is modulated at 70% duty cycle. This is associated with its higher matrix cell density and higher functional connectivity between the VA thalamus and transmodal cortical areas compared to other thalamic nuclei. **b** Categorization accuracy decreases at 70% duty cycle in a target-invariant manner, potentially driven by common interactions between the thalamus and the visual network (cyan). VA: ventral anterior thalamus; VP: ventral posterior thalamus; DA: dorsal anterior thalamus; DP: dorsal posterior thalamus; DC: duty cycle.

Being primarily nonspecific and matrix cell-rich, the VA thalamus likely interacts extensively with thick-tufted layer-5 (TTL5) pyramidal neurons in the cortex, potentially promoting large-scale cortical integration essential for conscious sensory processing^1,13,23,24,62^. This also supports theoretical frameworks such as thalamic gating mechanisms and dendritic integration theory, which suggest that matrix cells in higher-order thalamus modulate the threshold for conscious perception by influencing the bursting activity of TTL5 pyramidal neurons in the cortex^1,13,63–65^. These matrix cells, with their diffuse connectivity, are hypothesized to facilitate the entry of sensory content into consciousness, potentially by lowering the activation threshold of L5 pyramidal neurons^13,26^. Our LIFU intervention at high DC may have induced excitation of the matrix cells in the VA thalamus, possibly increasing dendritic integration in L5 pyramidal neurons and promoting widespread cortical activation that enhances perceptual sensitivity (further discussion on DC-dependence follows below).

The ventral anterior part of the thalamus (not to be confused with the ventral anterior thalamic nucleus) encompasses the intralaminar nuclei, which project to various cortical and subcortical regions^19,66^. Electrical stimulation of these nuclei via implanted electrodes has been shown to awaken non-human primates from anesthesia^18,67^ and aid in the recovery of consciousness in patients with neuropathological conditions^68,69^. Similarly, microinjection of nicotine or potassium channel-blocking antibodies into these nuclei has restored consciousness in rodents^70^. A previous LIFU study targeting these nuclei demonstrated improvements in patients with disorders of consciousness^36^. Our findings provide preliminary evidence that the intralaminar nuclei might also influence the contents of consciousness, in addition to modulating the global states of consciousness.

One important question is whether the increased sensitivity of conscious perception was due to an excitation of the VA thalamus. Previous empirical studies suggest that LIFU selectively activates excitatory vs. inhibitory neurons depending on the DC, thus exerting a general excitatory effect at high DCs (e.g., > 50%) and an inhibitory effect at low DCs (e.g., < 20%)^39,40,48,71–74^. Based on these observations and the subsequently established neuronal intramembrane cavitation excitation model^48^, we hypothesize that LIFU at 70% DC applied to the VA thalamus likely induced an excitatory effect in our study. This interpretation is also supported by relevant human neuroimaging studies. For example, Wu et al. demonstrated that higher pre-stimulus activity in the VA thalamus predicts greater sensitivity in conscious perception^12^. This is in line with our interpretation that 70% DC LIFU administered to the VA thalamus elevated its overall activity level, leading to heightened perceptual sensitivity. Furthermore, given the strong bidirectional excitatory connections^75^ and functional connectivity (Fig. 5a, d), LIFU applied to the VA thalamus may influence the activity of the mPFC. This is also supported by evidence that single-pulse electrical stimulation of the VA activates the mPFC^76^. Higher pre-stimulus mPFC activity also predicts greater perceptual sensitivity—a prediction intriguingly aligned with the previous findings^12^. Collectively, these observations suggest that 70% DC LIFU exerted an excitatory effect on the VA thalamus, which in turn induced co-activation of the mPFC, which could account for the observed increase in perceptual sensitivity.

Furthermore, we found that matrix cell richness, measured by the difference in CALB1 and PVALB expression levels (CP_T_)^22^, significantly correlated with enhancements in perceptual sensitivity. This relationship was specific to sensitivity and not observed for other outcomes, suggesting a distinct role for matrix cell-rich regions in modulating perceptual discrimination. Unlike region-based analyses that assume fixed anatomical targets, our approach incorporated individual beam profiles at the voxel level, accounting for variability in sonication sites and local cell composition. By linking behavioral effects to the actual sonication location and underlying cytoarchitecture, our findings offer more direct evidence that matrix cell density in the thalamus plays a causal role in conscious perception.

Unlike the increase in perceptual sensitivity with VA thalamus stimulation, categorization accuracy was reduced with all four thalamic targets at 70% DC (Fig. 6b). This target-invariant effect did not correlate with core-matrix cell composition, suggesting a distinct underlying mechanism. Given the excitatory nature of high-DC LIFU and the shared thalamovisual connectivity among the targeted regions (Fig. 5c), it is plausible that 70% DC sonication consistently co-activated the visual cortex during the LIFU-ON blocks^44^. Such co-activation may have broadly disrupted thalamocortical visual processing, affecting visual cortical networks that integrate inputs from multiple thalamic regions. This interpretation aligns with previous work demonstrating a negative correlation between the pre-stimulus activity of the visual network and categorization accuracy^12^.

It is possible that the target-invariant effects were confounded by auditory artifacts^77^. Sonication with 5% DC generated more noticeable noise from the transducer due to the higher pulse-averaged intensity than with 70% DC (14.4 W cm^-2^ for 5% DC vs. 1.03 W cm^-2^ for 70% DC), given a fixed temporal-average intensity. This led to a higher percentage of participants reporting awareness of the transducer noise at 5% DC (93% vs. 52% of participants, see Source Data). However, our results cannot be explained solely by auditory artifacts. First, when we included transducer noise awareness as a covariate in our statistical model, it showed no significant effect on behavioral outcomes (all *p*_FDR_ > 0.05; Supplementary Table 4). Second, if auditory artifacts were the primary driver of the perceptual changes, one should expect target-invariant effects, with a larger magnitude at 5% DC (at which transducer noise is more noticeable). Yet, the decrease in categorization accuracy occurred only at 70% DC (Table 1). Nonetheless, to better isolate and understand the effects of auditory artifacts, future research may include sham control by masking transducer noise or physically blocking the ultrasound beam.

While previous research demonstrated long-lasting LIFU effects ranging from hours to weeks^78,79^, we did not observe persistent changes as evidenced by the comparison between two baselines (Supplementary Fig. 3). This discrepancy may be attributed to two factors. First, the visual system, being highly adaptable and dynamic, might respond differently to LIFU compared to states like addiction, anxiety, or depression in which longer-term effects have been observed^74,78,80^. Second, the intensity of our LIFU modulation might have been insufficient to trigger long-term plasticity. To better understand the implications of LIFU for scientific study and therapeutic application, future investigations should assess the factors that influence the duration of sonication effects.

Methodologically, LIFU offers unique advantages for causal and mechanistic investigations of the neural substrates of consciousness in humans. Non-invasive techniques such as transcranial magnetic stimulation have been employed to transiently disrupt the contents of consciousness, for example, visual awareness^81–83^. However, these approaches are largely limited to superficial cortical regions and lack the spatial resolution necessary to target deep subcortical circuits. Other techniques such as deep brain stimulation and intracranial electrical stimulation applied to non-human primates^11,67^ or surgical patients^84–86^ have provided important insights into the neural mechanisms of conscious perception, but their invasiveness restricts clinical accessibility and broader applicability. By enabling non-invasive, focal neuromodulation of both cortical and deep subcortical structures with high spatiotemporal precision, LIFU provides a powerful new tool for dissecting the causal architecture of human consciousness.

Our work represents several refinements in the field of human LIFU research. First, in contrast to prior human studies that employed only one DC^87–89^, we incorporated two duty cycles (70% vs. 5%) into our experimental design and found DC-dependent effects. Second, while regional targeting subcortical regions has been demonstrated in rodents^40^, previous human LIFU investigations mainly targeted thalamus as a single target^35,74,87^. Our work extends these findings by systematically targeting various thalamic subregions with sub-centimeter precision. The average lateral deviation of the beam from the target was approximately 2 mm (Fig. 2b), smaller than beam width of about 5 mm. This was comparable to or better than those in previous works (3–9 mm)^87,90^. Therefore, our targeting approach suggests promising avenues for translational scientific investigations and clinical applications. Third, of relevance to future clinical applications, we mitigated the reliance on anatomical imaging (e.g., computed tomography or T1 MRI) by using a template rescaling method. We verified that this method meets the desired targeting accuracy in two ways: (1) using a separate MRI dataset, we confirmed small deviations of approximately 2 mm relative to the actual T1 image (Supplementary Fig. 4); (2) we observed that effect size increased as the beam-target lateral deviation decreased, which is a good indicator that our method effectively estimates the target location within an acceptable error range (Supplementary Fig. 11a). Our approach could expedite the experimental procedures and enable LIFU application in populations where anatomical scans are challenging, such as infants, individuals with implanted medical devices, or those with severe claustrophobia.

Our study has limitations. First, although the template rescaling approach described above allowed us to effectively approximate each participant’s thalamic coordinates, we did not create subject-specific acoustic models or apply skull-dependent phase corrections. The beam profiles therefore cannot capture the focal shift, peak-pressure drop, or off-axis energy leakage introduced by each participant’s skull morphology. This lack of individualized skull modeling represents a limitation that may have introduced unmodeled variability in beam propagation and targeting precision, potentially affecting both the magnitude and localization of the observed neuromodulatory effects. Likewise, while the transducer was positioned with an optical neuronavigation system, its final offset and tilt were not iteratively refined based on native MR or computed tomography images. This limitation further compromises targeting precision and should be addressed in future investigations through individualized anatomical imaging or automated transducer positioning protocols. Second, we did not perform functional imaging in the study participants; while our findings are consistent with previously suggested DC-dependent LIFU effects, we acknowledge that further investigation using concurrent functional imaging is necessary to conclusively prove the existence and directionality of neural activity and connectivity changes. Third, our simulations suggest significant energy loss (about 87%), mainly due to skull^53,54,57,91^, with potential additional attenuation by residual air pockets in hairs. Despite such loss, previous human studies using comparable or lower intensities have still demonstrated significant neuromodulatory effects^36,92–94^. In a recent rodent study, a temporal-average intensity as low as 0.0075 W cm^-2^ (roughly 100-fold lower than ours; about 10-fold lower after considering skull-induced attenuation) was sufficient to induce a detectable calcium signal^40^. Nonetheless, advanced approaches such as direct attenuation compensation can help mitigate these issues^91^. Currently, FDA intensity guidelines (e.g., derated spatial-peak temporal-average intensity *I_spta.3_* = 0.72 W cm^-2^) are based on standards for diagnostic and imaging ultrasound devices, and no specific guidelines exist for LIFU applications in the human brain. As studies in animal models have shown that higher intensities can yield stronger effects without causing tissue damage^40,95^, future work should advocate for updated FDA guidelines tailored to human brain LIFU.

In conclusion, our findings illuminate the causal roles of thalamic regions in conscious perception, particularly highlighting the unique contribution of the matrix cell-rich, transmodal-dominant VA thalamus in modulating visual discrimination sensitivity. These results provide insight into the relationship between cytoarchitectural and functional organization of the thalamus, and human conscious perception.

## METHODS

### Participants

The study protocol was approved by the Institutional Board Review of the University of Michigan Medical School. A total of 60 participants (age: 25.9 ± 6.3 years, mean ± SD; 38 females, 22 males) participated in the current experiment. All subjects provided written informed consent prior to the experiment and were compensated. All subjects were right-handed, did not have any hearing loss, were not colorblind, and had normal or corrected-to-normal vision. Although hair shaving was not required for this study, seven participants chose to shave their right temple, and they were given additional compensation in exchange. Participants were assigned to one of two equal-sized groups (duty cycle = 70% or 5%) by a random number generator, each group comprising 30 participants. Since the effect of LIFU neuromodulation is dependent on various factors, such as parameters and target regions, determining a prior effect size is challenging. However, we planned for a subject number of *n* = 30 per group, exceeding the mean sample size (*n* = 19) reported in the literature^96^. One subject who performed a different visual task was not included in data analysis. Three subjects with technical issues during targeting were excluded from data analysis. One subject did not complete the task due to technical issues in perceptual threshold determination. One subject was excluded because the hit rate during Baseline-1 was too low (< 15%). Thus, a total sample size was *n* = 54 (70% duty cycle: *n* = 27; 5% duty cycle: *n* = 27; see also Supplementary Fig. 1 for a comprehensive flow chart). Sex or gender analysis was not conducted because there were no sex or gender specific hypotheses regarding the influence of ultrasound neuromodulation on visual perception. Participants were blinded to the intervention conditions by providing identical procedural setups and device operations across both groups, ensuring that all participants did not know the specific duty cycle assignment.

### Low-intensity focused ultrasound devices and parameters

LIFU was administered using the BrainSonix BXPulsar 1002 System (BrainSonix Corporations, Sherman Oaks, CA, USA). This system includes a single transducer with a 61.5 mm diameter. The free-field focal depth is 80.7 mm, validated by Blatek Industries, Inc. (Boalsburg, PA, USA), following ASTM E1065/E1065M Standard guidelines (see Supplementary Table 2 for test data report). This transducer is mounted in a plastic housing and sealed with a thin polyethylene membrane. Free-field beam volume including length and width was verified by BrainSonix Corporations with a representative 80-mm transducer using a bilaminar membrane hydrophone with a 0.4 mm diameter active receive aperture (Supplementary Fig. 5). Brainsight Neuronavigation System (Rogue Research, Montreal, Quebec, Canada) was used for beam targeting.

Sonication parameters were as follows: fundamental frequency: 650 kHz; pulse repetition frequency: 10 Hz; pulse duration: 100 ms; duty cycle: 70% and 5% (yielding pulse width of 70 and 5 ms, respectively); sonication duration: 30 s; inter-sonication interval: 30 s; LIFU-ON epochs per block: 12 epochs; non-derated spatial-peak temporal average intensity (*I_spta.0_*): 1.02 W cm^-2^; derated spatial-peak temporal average intensity (*I_spta.3_*): 0.72 W cm^-2^; non-derated spatial-peak pulse-average intensity (*I_sppa.0_*): 1.03 W cm^-2^ (70% DC) and 14.4 W cm^-2^ (DC 5%); non-derated incident pressure (*P_r.0_*): 197 kPa (70% DC) and 710 kPa (5% DC); derated incident pressure (*P_r.3_*): 165 kPa (70% DC) and 619 kPa (5% DC). Since we used a fixed *I_spta.3_* for two duty cycles, we note that pulse-average intensity (*I_sppa_*) was higher at DC 5% condition. Given that the Mechanical Index (MI) is the derated pressure divided by the square root of the fundamental frequency (0.65 MHz), MI values were estimated as: 0.20 (70% DC) and 0.77 (5% DC), below the ITRUSST risk recommendations^58^. The derated values were calculated using the US FDA’s derating method, which assumes a uniform tissue attenuation rate of 0.3 dB cm^-1^ MHz^-1^. A comprehensive parameter report summary based on the ITRUSST reporting guidelines^97^ can be found in Supplementary Table 1.

### Transducer setup

Aquasonic Ultrasound Gel (Bio-Medical Instruments, Inc., Clinton Township, MI, USA) was applied to the right temple before positioning the transducer, ensuring that all hairs in the area were thoroughly coated with gel to enhance ultrasound transmission. The transducer was then positioned on the right temple and secured with two adjustable fabric bands: one band ran from the chin over the top of the head, and the other band passed from the forehead to the back of the head. Additional self-adhering bandages were used on top of the fabric bands to fine-tune the angle of the transducer for precise alignment. For post-hoc analysis, at the start and end of each LIFU-ON block, lateral deviation and coordinates of the ideal beam center (i.e., 80 mm away from the center of the transducer surface) and the head-transducer contact point were recorded from the neuronavigation system. Beam vector was then defined as a vector originating from the recorded head-transducer contact point, orienting toward the ideal beam center.

### Target locations

Following a gradient-based thalamic parcellation scheme^52^, the thalamus was coarsely divided into four regions: ventral anterior (VA), ventral posterior (VP), dorsal anterior (DA), dorsal posterior (DP). The center-of-mass coordinates for the four regions in the MNI space are as follows; Left VA: (-8 mm, -11 mm, 6 mm), Left VP: (-13 mm, -23 mm, 2 mm), Left DA: (-12 mm, -23 mm, 12 mm), Left DP: (-16 mm, -31 mm, 2 mm).

### Head size measurement

We used a digital outside-diameter caliper to measure head size in three dimensions: width, depth, and side-to-top distance (Fig. 1b). Head width was defined as the distance between the left and right supratragic notches, which are indentations in the ear cartilage right above the tragus. Head depth was measured from the most posterior (back) point to the most anterior (front) point of the head. The side-to-top distance was defined as the measurement from the one supratragic notch to the highest top point of the head.

### Template rescaling method

We rescaled the ICBM 2009c Nonlinear Asymmetric template of the MNI152 linear template to match each participant’s head size based on the three measured dimensions. The head height, defined as the height of a triangle formed by the top of the head, the supratragic notch, and the center of the head at the plane of the supratragic notch, was calculated using the formula:

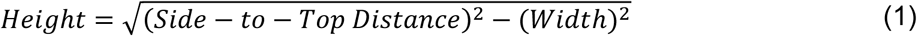

The default head size of the MNI template were as follows: width: 16 cm, depth: 21 cm, and height: 15 cm. We then calculated the size ratio between the MNI template and the participant’s head dimensions. The NIFTI image of the MNI template was resliced using B-spline interpolation to adjust to the participant’s head size.

We performed a post-hoc analysis with a separate MRI dataset (*n* = 13) to assess the accuracy of our template rescaling method. For each participant, we obtained an anatomical T1 image and measured head size (width, depth, and side-to-top distance). We then created a pseudo-anatomical rescaled template as described above. The actual T1 and the rescaled template images were aligned according to the anterior and posterior commissures, segmented, normalized to MNI space, and used to generate inverse deformation fields. We applied the inverse deformation fields to map four target MNI coordinates into each image’s native space and computed the Euclidean distances between the resulting coordinates for each thalamic region.

This distance provided a quantitative measure of how closely the rescaled template matched each participant’s true anatomy at the thalamic targets.

### Simulation of ultrasound beam

Simulation of ultrasound intensity and thermal profiles was performed using a two-step segmentation and modeling approach. We initially segmented the pseudo-anatomical image into scalp, skull, and brain employing the Complete Head Anatomy Reconstruction Method (CHARM)^98^. For a more accurate segmentation, we used a fat-suppression method (https://github.com/ProteusMRIgHIFU/EnhanceTUS_Zadeh_JNE)^99^. For cases where participant-specific ultrasound trajectories extended beyond the headspace due to anatomical variability, the label assignment of scalp was manually extended outward by 1 cm, and the mesh was recalculated with CHARM, ensuring that all recorded head-transducer contact points lie within the adjusted headspace.

Subsequently, acoustic property assignments and ultrasound beam simulations were performed using an open-source software named BabelBrain^100^. Simplified masks based on the CHARM- generated MRI segmentations were utilized, with trabecular bone proportion set at 80%. These masks assigned homogenized acoustic properties, including density, longitudinal and shear speeds of sound, and attenuation, derived from established literature values^100^. Simulations were conducted up to 100 mm below the transducer with a spatial resolution of six points-per-wavelength.

The tissue attenuation ratio was determined by how much acoustic energy is lost when ultrasound propagates through tissue compared to a water reference (exported as “RatioLosses” variable in BabelBrain software). In short, the software selects the plane at which the maximum tissue pressure occurs. It then computes the acoustic energy in this plane as the sum over all voxels of the squared pressure amplitude. Similarly, the code processes the free-field pressure amplitude data at the same plane. The tissue attenuation ratio is then defined as the ratio of the tissue acoustic energy to the water acoustic energy at that specific plane.

Thermal effects resulting from ultrasound exposure were estimated using the Bio Heat Transfer Equation (BHTE). Baseline temperature for thermal modeling was set at 37 °C. During the simulation, BabelBrain software calculates the temperature evolution in the tissue (i.e., skin, skull, brain, and target point) both during ON and OFF epochs, which are then concatenated, yielding an array of 4 × 72,000 size. Maximal temperature was determined by the maximal value within this array.

The simulated relative intensity values exported from BabelBrain software were normalized to MNI space using the forward deformation fields that MNI-normalized the corresponding pseudo-anatomical image and then 3D-interpolated using ‘interp3’ function in MATLAB with the refinement factor of 4. The individual beam trajectory images in Supplementary Fig. 8-10 were also MNI-normalized and interpolated. The spatial coverage of each beam was determined as the percentage of “beam-present” voxels (defined as voxels exhibiting more than 50% of the relative intensity^101^) that fell within specific thalamic (or other non-thalamic) regions. For example, the sum of beam coverage values across 10 regions (eight thalamic and two non-thalamic regions) equals 100%.

### Overall experimental procedure

The experiment was conducted in a single session lasting approximately two hours, where the main task took approximately 1.5 hours (Fig. 1a). First, we measured the participant’s head size (details above), followed by task instructions. The participant then completed a brief practice run consisting of 30 trials (a shorter version of staircase procedure; see below) while the experimenter generated the pseudo-anatomical image. Next, the ultrasound transducer was positioned, and participants underwent the image contrast staircase procedure involving 60 trials. After confirming the convergence of contrast value (see below), participants then engaged in the main task, which consisted of six blocks, each containing 100 trials and lasting approximately 11.5 minutes (Fig. 1a). Each block featured four trials per each real image (80 real trials in total) and five trials per each scrambled image (20 scrambled trials in total; see below for details in visual stimuli). Participants were unaware of the fact that scrambled images would appear.

During the inner four blocks, LIFU was administered toward one of four left thalamic areas in a pre-assigned, pseudo-randomized order (determined by a random number generator), counter- balanced across the participants. Within one participant, only one duty cycle (either 70% or 5%) was applied throughout the four LIFU-ON blocks. Participants were allowed to rest between blocks (3.8 ± 2.4 min; mean ± SD; see Source Data), during which the transducer was readjusted. We also measured the eye-to-screen distance and readjusted the position of the laptop if necessary (see below). After the transducer was successfully repositioned and the participants were willing to proceed, the visual task was self-initiated by participants and LIFU administration was manually triggered by the experimenter after the task onset.

### Visual stimuli

Visual images encompassed four categories: faces, houses, human-made objects, and animals (Fig. 1g), which were used in a previous study^8^. The images were sourced from publicly available labeled photographs or the Psychological Image Collection at Stirling (PICS; https://pics.stir.ac.uk). Each category included five distinct images, resulting in a total of 20 unique real images. All images were converted to grayscale and resized to 300 × 300 pixels. The pixel intensities were normalized by subtracting the mean and dividing by the standard deviation. Subsequently, the images were processed with a 2D Gaussian smoothing filter, using a standard deviation of 1.5 pixels and a 7 × 7 pixel kernel. Scrambled images that maintain low-level features were generated for each category by randomizing the phase of the 2D Fourier transformation for one image from each category.

The contrast of an image was defined as:

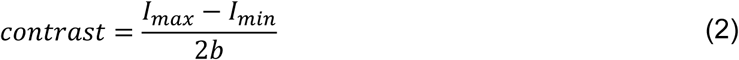

where *I_max_* and *I_min_* are the intensity values of lightest and darkest pixels in the image, respectively (in the range of 0 to 255). To ensure a smooth transition to the background (set as gray, intensity = 127), the edges of all images were gradually blended by applying a Gaussian window with a standard deviation of 0.2 to the image intensity.

The stimuli were displayed on a 14-inch laptop (HP ProBook 440 G4) with a 60-Hz refresh rate LCD screen, set to the maximum screen brightness. The viewing distance between the screen and the participants’ eyes was maintained at 65.8 ± 5.7 cm (mean ± SD). During the experiment, participants were asked to maintain constant eye-monitor distance. We did not adopt masking because masking is known to influence the visual information processing^9^.

### Image contrast staircase procedure

Before the main task, participants completed the QUEST adaptive staircase procedure^8^ to determine the contrast value that would yield a 50% recognition rate (i.e., the rate of “YES” response to Question-2). The QUEST procedure was similar to the main task but with shorter timing parameters and without scrambled images. The inter-trial interval was 0.75 seconds, and the delay period between the stimulus and the first question was 2 seconds. The threshold contrast was identified through a QUEST process consisting of 60 trials. Each trial was randomly assigned to one of three sets, and the staircase procedure was conducted separately for each set. Task performance was deemed acceptable if the three sets successfully converged on a specific contrast value. The staircase procedure was repeated until the convergence of the three curves was confirmed. Rather than adjusting the contrast of each image individually, the staircasing aimed for 50% recognition across all images.

### Trial structure

Each trial began with a fixation cross displayed on a gray background for a randomly determined duration between 3 and 6 seconds, following an exponential distribution to prevent participants from predicting stimulus onset (Fig. 1e). The stimulus image was then shown behind the fixation cross for 8 frames (67 ms), with the image intensity linearly increasing across frames. After the stimulus disappeared, the fixation cross remained on the screen for an additional 2 seconds. Each trial concluded with two sequential questions about the stimulus, each displayed for up to 3 seconds.

Question-1 asked participants to categorize the image as a face, house, object, or animal. Participants were instructed to guess the category even if they did not consciously recognize the object (four alternative forced-choice). Question-2 assessed their subjective experience, asking whether they had a "meaningful visual experience" of the object stimulus ("YES" or "NO"). Before the practice run, participants were told that a "meaningful" stimulus was defined as something that makes sense in the real world, as opposed to random noise or meaningless shapes. They were instructed to respond "YES" even if they recognized only part of an object. Participants provided their answers by pressing buttons on a 4-key external USB keyboard placed on their lap. If participants did not respond to Question-1, it was recorded as incorrect, and for Question-2, a lack of response was recorded as “NO.”

### Signal detection theory analysis

We employed signal detection theory (SDT) to examine the perceptual outcomes, focusing on sensitivity and criterion^9^. Sensitivity measures the ability to distinguish between real images and scrambled images. We calculated parametric sensitivity metric *d’* through standard SDT analysis defined as follows:

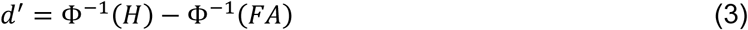

where Φ^−1^ is the inverse normal cumulative distribution function, *H* is the hit rate, and *FA* is the false alarm rate, respectively. High *d’* indicates better discrimination between real and scrambled images.

Decision bias refers to the inclination to certain response, regardless of whether the stimulus is real or scrambled. We assessed decision bias using parametric metric criterion *c* defined below.

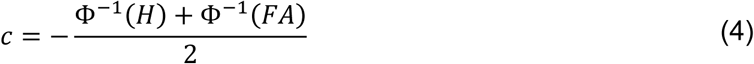

High *c* values indicate the tendency to say “NO” to Question-2, indicating a more conservative object recognition.

### Statistics

To ensure stabilization of task performance, the first ten trials of each block were omitted from the analysis. Due to technical issues, a few participants failed to complete specific LIFU-ON blocks, which were consequently excluded from the analysis. The numbers of affected participants for each condition are as follows: 70% DC (VA: *n* = 1; VP: *n* = 1) and 5% DC (VA: *n* = 2; VP: *n* = 2; DA: *n* = 1). For one subject, the lateral deviation during VP sonication was not recorded. The coordinates for head-transducer contact point and ideal beam center were not recorded for a few participants, resulting in their omission from simulation analysis. The numbers of affected participants were as follows; VA: *n* = 8; VP: *n* = 9; DA: *n* = 8; DP: *n* = 8. Because *d’* and *c* are ill-defined at perfect performance (i.e., Φ^−1^(𝐻𝐻) = ∞ at *H* = 1)^102,103^, we omitted such blocks with from the statistical analyses of *d’* and *c*.

To address potential confounds, we included six nuisance variables in the analysis. Subject-specific factors correspond to age, sex, and the awareness of transducer noise during LIFU-ON blocks (coded as 0 for unaware, 0.5 for unsure, and 1 for aware). Block-specific factors such as lateral deviation, sonication order, and tissue attenuation ratio (in dB) were also incorporated. Missing data for noise awareness was imputed using modal category (noise awareness: 1). Missing or baseline values of lateral deviation and attenuation ratio were imputed using group means (lateral deviation: 2.251 mm; attenuation ratio: -8.7812 dB). Given no significant differences between Baseline-1 and Baseline-2 across all measures (Supplementary Fig. 3), we averaged these into a single baseline measure. No imputation was performed for the main perceptual outcome data.

We implemented a sequential analytical approach beginning with omnibus testing using linear mixed-effects (LME) models. This approach was chosen because LME models can directly accommodate covariates, handle non-normal data distributions, and properly account for missing outcome data through restricted maximum likelihood estimation. Our statistical implementation involved fitting a single linear mixed-effects model specified as:

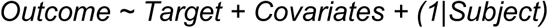

where *Target* includes baseline and four LIFU regions (five categories total) and *Covariates* encompass the six nuisance variables described above. For each perceptual outcome, we first conducted an omnibus F-test to determine whether there were any differences among the five experimental conditions using MATLAB’s ’fitlme’ and ’anova’ functions with Satterthwaite approximation.

Only outcomes that demonstrated significant omnibus effects after FDR correction proceeded to post-hoc analysis. For significant outcomes, we conducted pairwise comparisons between each LIFU condition and baseline using linear contrasts, with additional FDR correction applied within this family of contrasts at *α* = 0.05. This approach ensures that each dataset undergoes a single, unified analytical pathway while controlling family-wise error rate at the outcome level. For the entire FDR correction steps, we used Benjamini-Hochberg procedure using MATLAB’s ‘mafdr’ function. Full statistics of main and supplementary analyses are included in Source Data.

### Thalamic Cytoarchitecture Analysis

To examine the relationship between thalamic cytoarchitecture and perceptual outcomes, we analyzed the core-matrix cell composition of stimulated regions. Thalamic cell types were inferred from the mRNA expression levels of CALB1 and PVALB, as provided by the Allen Human Brain Atlas^59^. The probes CUST_11451_PI416261804 and A_23_P17844 were used to estimate PVALB expression. For estimating CALB1 expression, CUST_140_PI416408490, CUST_16773_PI416261804, and A_23_P43197 were used. The CP_T_ metric was computed as the weighted difference between CALB1 and PVALB expression levels^22^. Higher CP_T_ values reflected greater CALB1 expression (i.e., matrix cell-rich regions), while lower values reflected greater PVALB expression (i.e., core cell-rich regions).

Using the thalamic CP_T_ map provided by James M. Shine’s Lab (‘calb_minus_pvalb.nii’ file in https://github.com/macshine/corematrix), we calculated the mean CP_T_ value within the beam- present voxels for each LIFU-ON block, defined as voxels exceeding 50% relative intensity based on acoustic simulations. These CP_T_ values were then correlated with baseline-subtracted changes in the three perceptual outcomes that survived omnibus testing.

### Human Connectome Project Dataset

The dataset was obtained from the S1200 Release of the WU-Minn Human Connectome Project (HCP) database, which has been extensively described in prior studies^60^. The participants were healthy young adults aged 22 to 37 years. Participants who had completed two sessions of resting-state fMRI scans (Rest1 and Rest2) were included in our analysis, resulting in *n* = 1009. The data were collected using a customized Siemens 3T MR scanner (Skyra system) with multiband EPI. Each scanning session comprised two sequences with opposite phase encoding directions (left-to-right and right-to-left), each lasting 14 minutes and 33 seconds. The sequences were acquired with a repetition time (TR) of 720 ms, an echo time (TE) of 33.1 ms, and a voxel size of 2 mm isotropic. To maximize data quality and minimize bias from phase encoding direction, the sequences from each session were combined, resulting in a total of 29 minutes and 6 seconds. The denoised volumetric data, preprocessed through ICA-FIX, were accessed from the online HCP database. Further details on the resting-state fMRI data collection and preprocessing are available in previous publications^60^. We employed standard preprocessing procedure, including resampling to a 3 × 3 × 3 mm resolution, band-pass filtering within the 0.01–0.1 Hz frequency range, spatial smoothing with a 6-mm Full Width at Half Maximum isotropic Gaussian kernel, and temporal normalization to attain zero mean and unit variance.

### Network-level analysis

We parcellated the cortex into 400 regions of interest (ROIs) based on a well-established parcellation scheme, assigning each ROI to one of seven canonical cortical networks: visual, somatomotor, dorsal attention, ventral attention, limbic, frontoparietal, and default-mode networks^61,104^. Additionally, we used the Subcortical Atlas^52^ to extract BOLD signal time courses from 14 distinct thalamic ROIs, including bilateral ventral anterior inferior (VAi), ventral anterior superior (VAs), ventral posterior lateral (VPl), ventral posterior medial (VPm), dorsal anterior medial (DAm), dorsal anterior lateral (DAl), and dorsal posterior (DP) regions. For analysis purposes, we averaged bilateral and related subdivisions into four primary thalamic regions with equal weights: VA (left and right VAi and VAs), VP (left and right VPl and VPm), DA (left and right DAm and DAl), and DP (left and right DP). For each resting-state session (Rest1 and Rest2), we computed the functional connectivity between each of these four thalamic regions and the 400 cortical ROIs using Fisher-*z* transformed Pearson correlation coefficients. Subsequently, for each session and thalamic region, we converted the correlation values into percentile ranks across all 400 cortical ROIs. We then averaged these percentile ranks within each of the seven canonical networks and across both resting-state sessions to derive network-level thalamocortical connectivity measures. Finally, we calculated the average rank scores for unimodal (visual and somatomotor) and transmodal (frontoparietal and default-mode) networks, visualizing these results in a 2D plot to illustrate relative connectivity patterns.

### Voxel-level analysis

As described above, mean time courses were extracted from bilateral VA, VP, DA, and DP thalamus, serving as seeds for voxel-level functional connectivity analysis. Seed-based maps (Fisher-*z* transformed Pearson correlation) were generated for each participant and region. Group-level *z*-score maps were generated by standardizing individual connectivity values (subtracting the mean and dividing by the standard deviation across participants). Within the cortex, we retained the top 10% of cortical voxels with the highest connectivity. This analysis was performed independently for Rest1 and Rest2 sessions, and their overlap was used to ensure consistency. Region-specific and region-invariant clusters of high-connectivity cortical voxels were then identified.

### Software

The staircase and main task were based on a custom-modified version of the publicly-available code for the near-threshold behavioral paradigm (https://github.com/BiyuHeLab/NatCommun_Levinson2021/) utilizing the Psychophysics Toolbox v3.0.19.4 (http://psychtoolbox.org/) executed on MATLAB R2023b^8^. Neuronagivation processes including transducer targeting and coordinate recording were conducted on proprietary Brainsight software (https://www.rogue-research.com/downloads/). Template rescaling and its post-hoc validation were performed with custom-built code and SPM12 (https://www.fil.ion.ucl.ac.uk/spm) both executed on MATLAB R2024a. Head segmentation with CHARM was conducted from the SimNIBS v4.1.0 (https://simnibs.github.io/simnibs/build/html/index.html). Ultrasound beam simulation was performed on BabelBrain v0.4.3 (https://proteusmrighifu.github.io/BabelBrain/). All subsequent analyses involving volumetric beam intensity data (e.g., MNI normalization, average beam intensity calculation, and CP_T_ calculation) were performed with custom scripts utilizing SPM12 on MATLAB R2024b. The 3D renderings of brain and beam overlaps were performed using MRIcroGL v1.2.20220720 (https://www.nitrc.org/projects/mricrogl). For fMRI data preprocessing and voxel-level connectivity analysis, we employed the AFNI software suite (linux_ubuntu_16_64; http://afni.nimh.nih.gov/). All other analyses (e.g., outcome calculation, statistical tests, and network-level connectivity analysis) and visualizations were conducted on MATLAB R2024b.

## DATA AVAILABILITY

Source data are provided with this paper. The HCP dataset is available from online repository (https://www.humanconnectome.org/). The thalamic CP_T_ map can be downloaded from Github repository (https://github.com/macshine/corematrix).

## CODE AVAILABILITY

Original code for near-threshold paradigm is available at Github (https://github.com/BiyuHeLab/NatCommun_Levinson2021/)^8^. Custom-built code for template rescaling is available at Github (https://github.com/janghw4/template_rescaling).

## ACKNOWLEDGEMENTS

We express our sincere gratitude to our study coordinators, Amy McKinney and Aaron Ellis, for their invaluable contributions to participant recruitment, meticulous scheduling, and ensuring the smooth execution of this study. We sincerely appreciate Drs. Mark E. Schafer and Alexander Bystritsky for providing free-field beam characteristics and Dr. Kelli A. Sullivan for thoughtful comments on skull-induced energy loss. This work was supported by the National Institute of General Medical Sciences of the National Institutes of Health grants R01GM103894 (to A.G.H. and Z.H.). The content is solely the responsibility of the author and does not necessarily represent the official views of the National Institutes of Health.

## AUTHOR CONTRIBUTIONS STATEMENT

H.J. and Z.H. designed the study and set up the experimental apparatus. H.J. devised the template rescaling method and developed the code. H.J., P.F., and Z.H. conducted the experiments. H.J. analyzed the data, prepared the figures, and drafted the manuscript. Z.H. co-analyzed the data. Z.H., G.A.M. and A.G.H. interpreted the data and edited the manuscript.

