## Supplementary for "Role of Thalamus in Human Conscious Perception Revealed by Low-Intensity Focused Ultrasound Neuromodulation"

### Perceptual outcomes at Baseline-1

We evaluated the perceptual outcome of the first block of the main task (Baseline-1) in which LIFU is not yet applied. The recognition rate (i.e., hit rate) was 53.3 ± 16.1% (mean ± SD), not significantly different from the intended target of 50% ($p = 0.09$, Wilcoxon signed-rank test, two-sided; Supplementary Fig. 2a). The recognition rate for scrambled images (i.e., false alarm rate) was 21.8 ± 17.3%, significantly higher than zero ($p < 0.0001$, Wilcoxon signed-rank test, two-sided) but lower than hit rate ($p < 0.0001$, two-sided Wilcoxon signed-rank test).

The sensitivity metrics showed $d' = 0.96 \pm 0.67$, significantly different from zero ($p < 0.0001$, Wilcoxon signed-rank test, two-sided; Supplementary Fig. 2b). All participants exceeded $d' = 0$, confirming that they could distinguish between real and scrambled images to some extent. Criterion $c$ (0.34 ± 0.38) was significantly greater than zero ($p < 0.0001$, Wilcoxon signed-rank test, two-sided; Supplementary Fig. 2c). This indicates a bias toward responding "NO" in Question-2, consistent with previous findings[1].

Categorization accuracy for real images was 84.9 ± 16.5% for recognized images, significantly higher than 47.7 ± 18.0% for unrecognized ones ($p < 0.0001$, Wilcoxon signed-rank test, two-sided; Supplementary Fig. 2d). Even unrecognized images had accuracy above chance level (25%), suggesting unconscious processing ($p < 0.0001$, Wilcoxon signed-rank test, two-sided; Supplementary Fig. 2d, red violin). Scrambled images showed a similar pattern, with recognized images at 72.1 ± 31.6% accuracy and unrecognized ones at 35.5 ± 13.9%, both lower than for real images (Supplementary Fig. 2e). Unrecognized scrambled images still had accuracy above chance ($p < 0.0001$, Wilcoxon signed-rank test, two-sided; Supplementary Fig. 2e, red violin), indicating that low-level features contributed to unconscious visual processing[2]. See Source Data for full statistics.

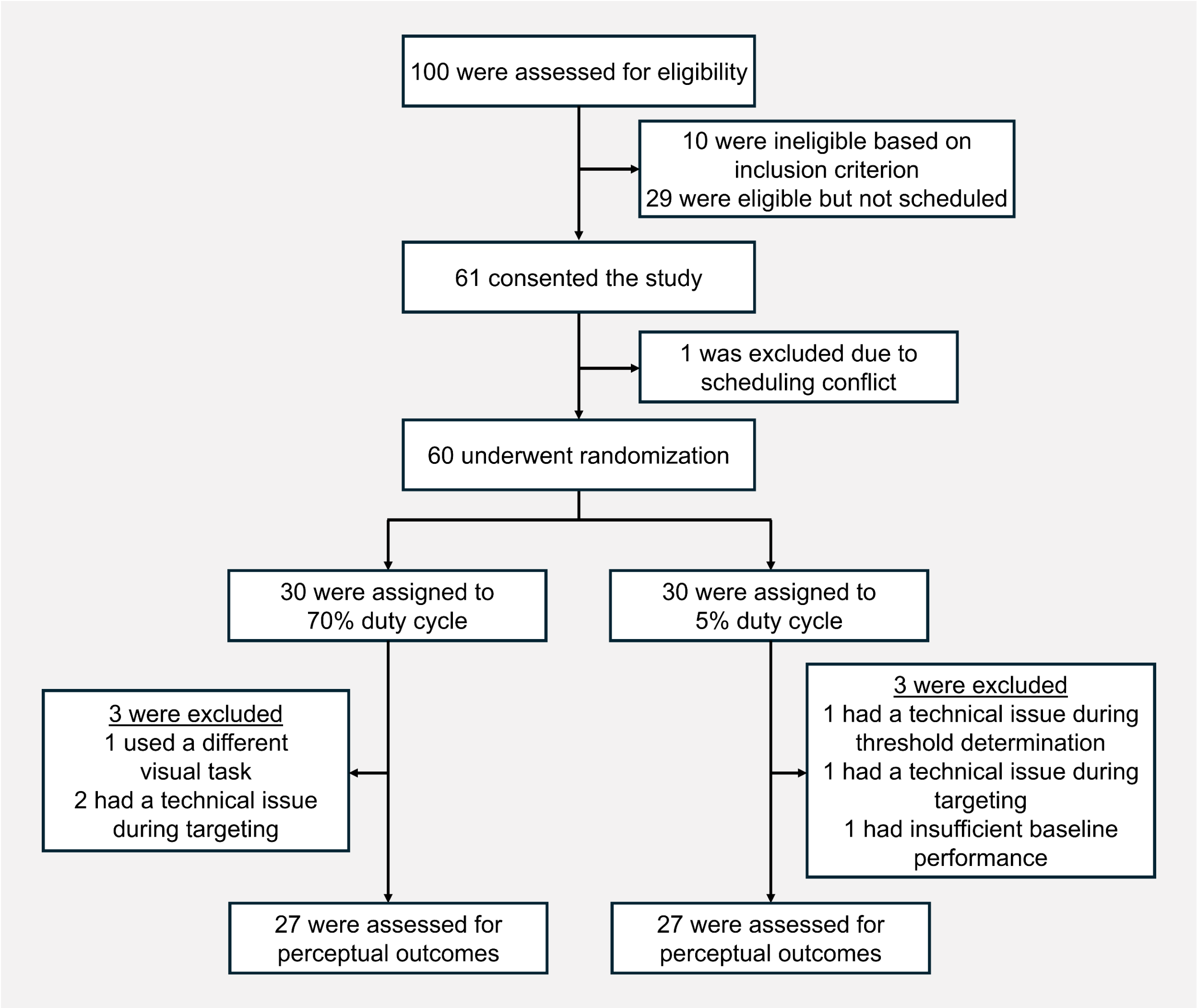


**Supplementary Fig. 1: Flow diagram of participant enrollment, randomization, and analysis.** Sixty participants were randomized into two groups: 30 assigned to the 70% duty cycle and 30 to the 5% duty cycle. Three participants from each group were excluded due to various experimental issues, leaving 27 participants per group for perceptual outcome assessment.

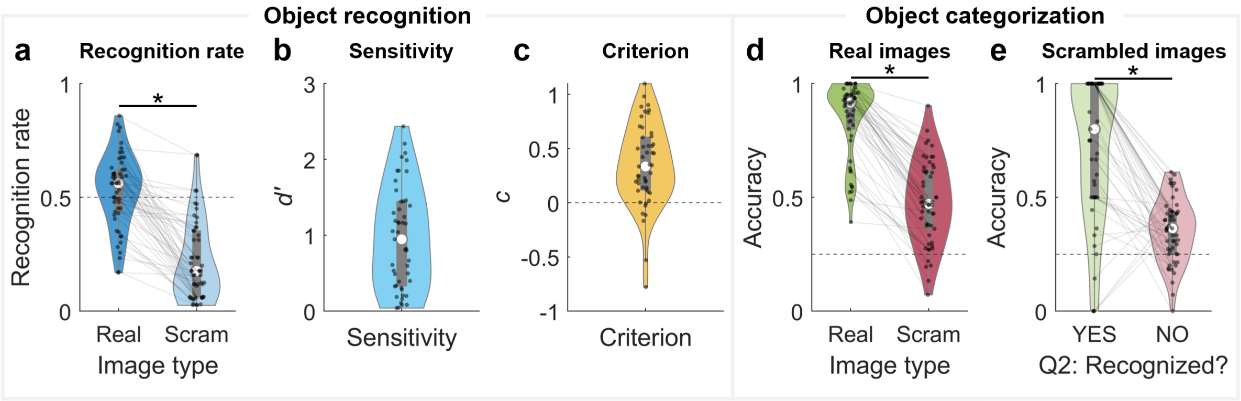


**Supplementary Fig. 2: Perceptual outcomes at Baseline-1. a** Recognition rate for real and scrambled (denoted "Scram") images, **b** Sensitivity *d′*, **c** Criterion *c*, **d** categorization accuracy of recognized vs. unrecognized real image trials, and **e** categorization accuracy of recognized vs. unrecognized scrambled image trials. Asterisks indicate statistical significance ($p < 0.05$, Wilcoxon signed-rank test, two-sided). Dashed lines are reference values for each measure: recognition rate of 0.5 (a), no bias of $c = 0$ (c), and chance accuracy of 0.25 (d, e). Gray boxes indicate interquartile ranges. Median values are marked by white circles. Sample sizes are as follows: (a,d) $n = 54$; (b,c) $n = 49$; (e) $n = 49$ (YES), $n = 54$ (NO). Statistics and source data are provided as a Source Data file.

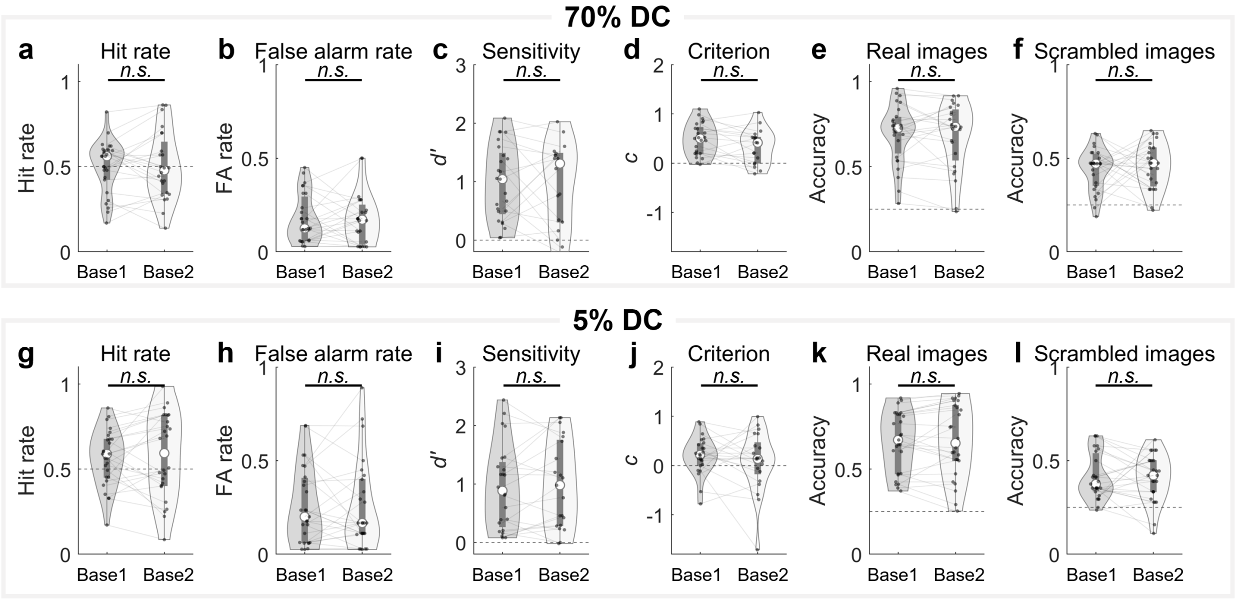


**Supplementary Fig. 3: Comparison of perceptual outcomes at baselines for two duty cycles (DC). a-f** Perceptual outcomes of 70% DC: **a** hit rate, **b** false alarm rate, **c** sensitivity *d′*, **d** criterion *c*, **e** categorization accuracy for real images, and **f** categorization accuracy for scrambled images. **g-l** Perceptual outcomes of 5% DC: **g** hit rate, **h** false alarm rate, **i** sensitivity *d′*, **j** criterion *c*, **k** categorization accuracy for real images, and **l** categorization accuracy for scrambled images. The *n.s.* denotes no significant difference ($p > 0.05$, Wilcoxon signed-rank test, two-sided). Dashed lines are reference values for each measure: hit rate of 0.5 (a, g), no discrimination of $d' = 0$ (c, i), no bias of $c = 0$ (d, j), and chance accuracy of 0.25 (e, f, k, l). Gray boxes indicate interquartile ranges. Median values are marked by white circles. Sample sizes are as follows: (a,b,e,f) Baseline-1: $n = 27$; Baseline-2: $n = 24$; (c,d) Baseline-1: $n = 25$; Baseline-2: $n = 18$; (g,h,k,l) Baseline-1: $n = 27$; Baseline-2: $n = 26$; (i,j) Baseline-1: $n = 24$; Baseline-2: $n = 21$. Statistics and source data are provided as a Source Data file.

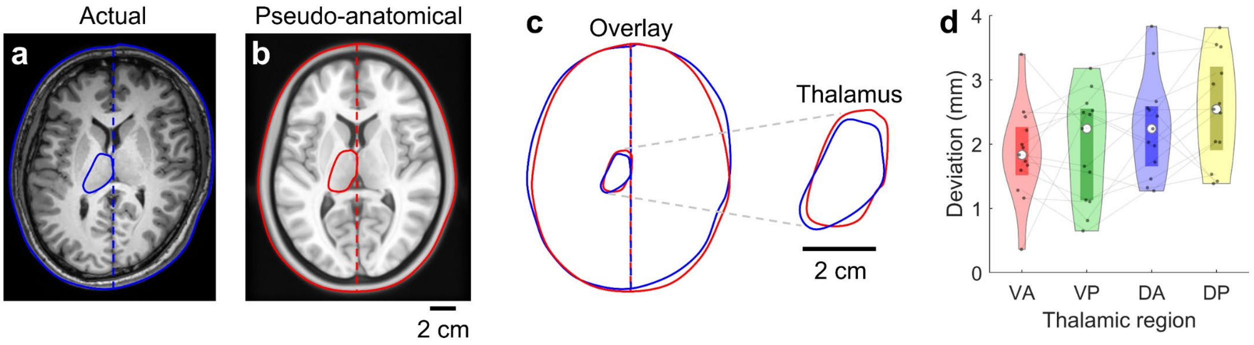


**Supplementary Fig. 4: Validation of template rescaling method. a** An example axial view of the actual anatomical T1 image with the scalp and left thalamus contour outlined in blue. **b** Equivalent axial view of the pseudo-anatomical rescaled MNI image with the scalp and left thalamus contour outlined in red. **c** Overlay of the scalp and thalamic contours from the actual and pseudo-anatomical images, qualitatively demonstrating the alignment and deviation between the two methods. The zoomed-in view highlights the left thalamus. **d** Quantification of the deviation between actual and pseudo-anatomical images for each thalamic region. Boxes indicate interquartile ranges ($n$ = 13). Median values are marked by white circles. VA: ventral anterior thalamus; VP: ventral posterior thalamus; DA: dorsal anterior thalamus; DP: dorsal posterior thalamus. Source data is provided as a Source Data file.

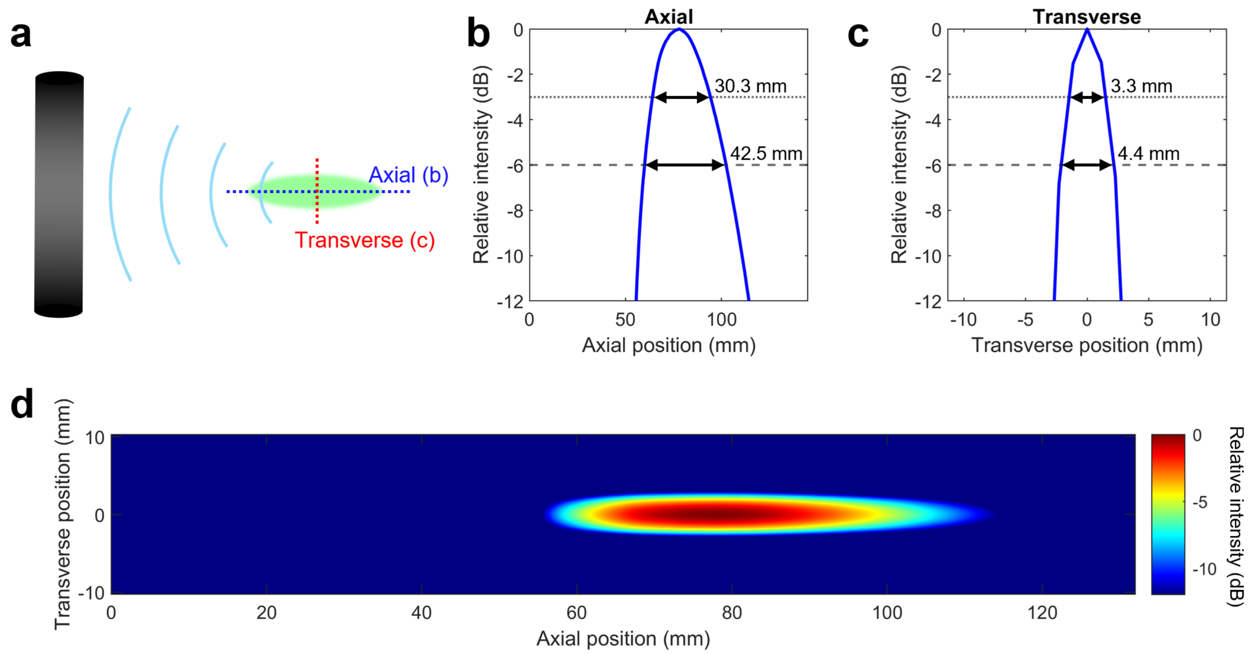


**Supplementary Fig. 5: Free-field acoustic profile of the transducer. a** Schematic illustration of intensity measurements at planes relative to the transducer. **b, c** Relative beam intensity across **b** axial and **c** transverse directions. Dotted and dashed lines indicate -3 dB and -6 dB (equivalent to 50% and 25% of the initial intensity). **d** Reconstructed two-dimensional (2D) axial-transverse intensity map of the acoustic beam profile. This data was provided by BrainSonix Corporations. Source data is provided as a Source Data file.

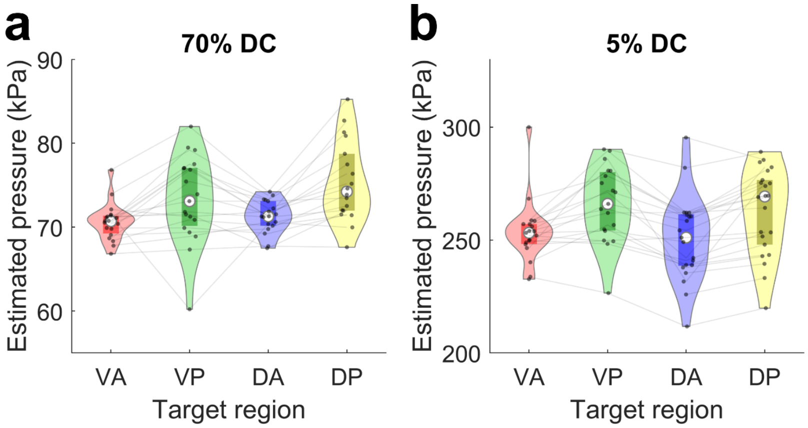


**Supplementary Fig. 6: Estimated tissue pressure.** Target-specific pressure values for **a** 70% and **b** 5% duty cycles (DCs). Boxes indicate interquartile ranges. For panel a, VA: *n* = 19; VP: *n* = 19; DA: *n* = 18; DP: *n* = 18. For panel b, VA: *n* = 19; VP: *n* = 19; DA: *n* = 21; DP: *n* = 22. Median values are marked by white circles. VA: ventral anterior thalamus; VP: ventral posterior thalamus; DA: dorsal anterior thalamus; DP: dorsal posterior thalamus. Source data is provided as a Source Data file.

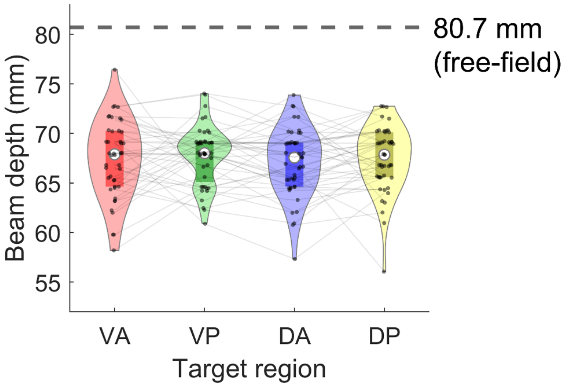


**Supplementary Fig. 7: Simulated beam depth across four targeted thalamic regions.** The beam depth is defined as the Euclidean distance from the head-transducer contact point to the location of maximal simulated intensity. Dashed line indicates focal depths measured in free field (80.7 mm). Boxes indicate interquartile ranges (VA: $n$ = 45; VP: $n$ = 44; DA: $n$ = 46; DP: $n$ = 46). Median values are marked by white circles. VA: ventral anterior thalamus; VP: ventral posterior thalamus; DA: dorsal anterior thalamus; DP: dorsal posterior thalamus. Source data is provided as a Source Data file.

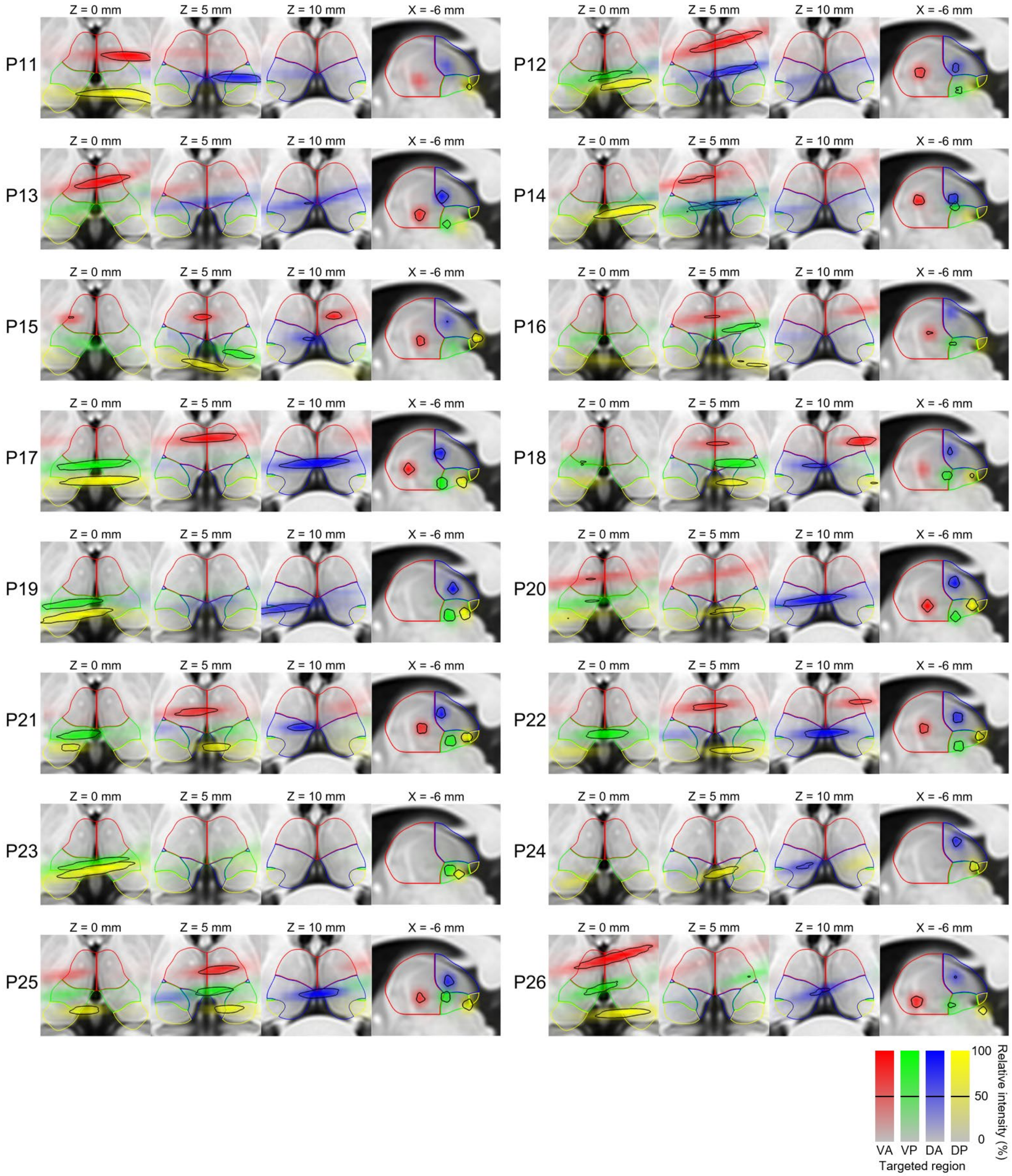


**Supplementary Fig. 8: Individual-level mosaic views of beam trajectories for participants P11 to P26.** Black contours indicate 50% (-3 dB) relative intensity. Color coding: VA (red), VP (green), DA (blue), and DP (yellow) thalamus. Some beams were not drawn due to reasons as follows: unrecorded head-transducer contact coordinates: entire beams of P02 – P09 and VP of P11; technical issues during certain blocks: VA of P19, VA and DA of P23, and VA and VP of P24. VA: ventral anterior thalamus; VP: ventral posterior thalamus; DA: dorsal anterior thalamus; DP: dorsal posterior thalamus.

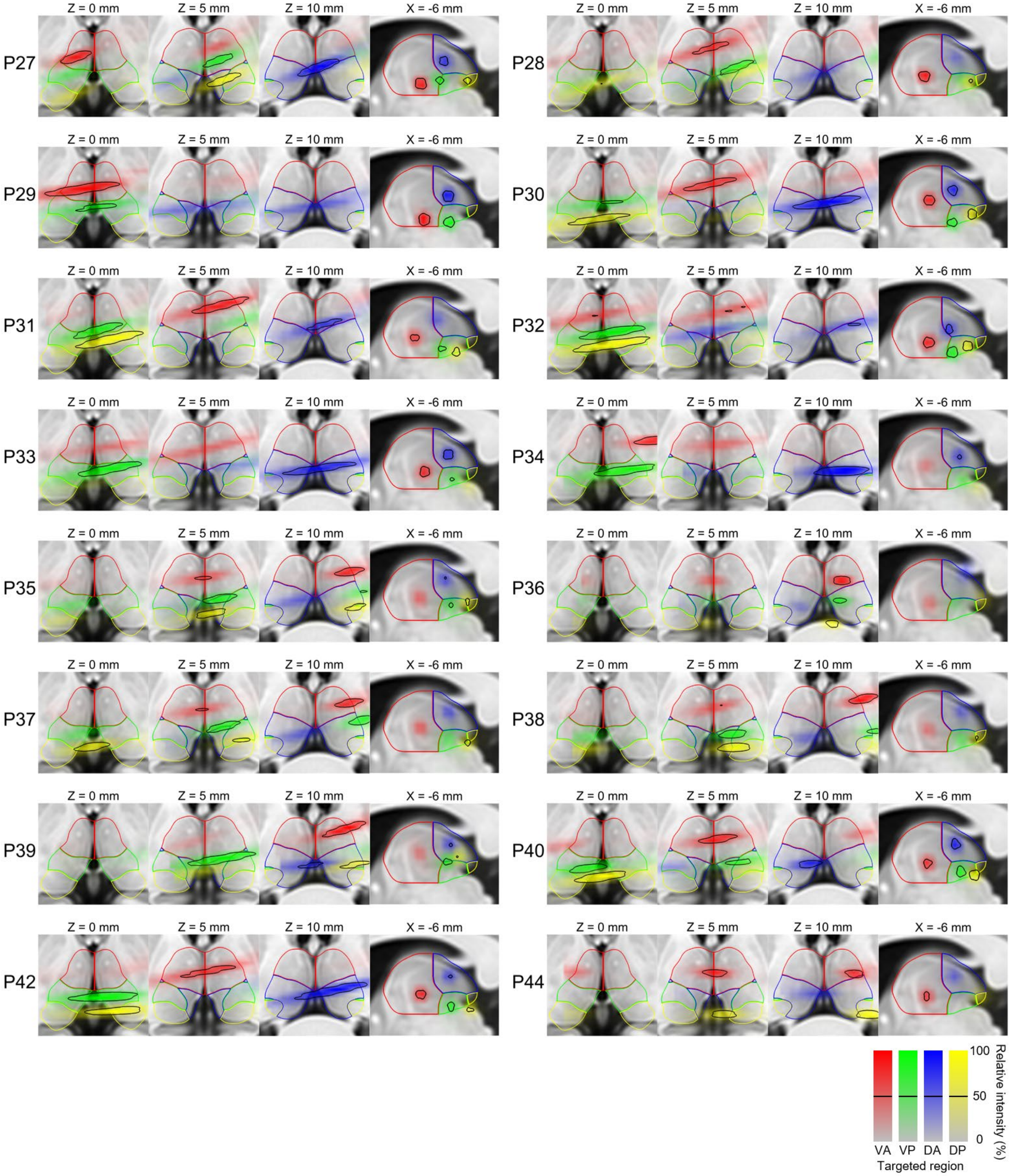


**Supplementary Fig. 9: Individual-level mosaic views of beam trajectories for participants P27 to P44.** Black contours indicate 50% (-3 dB) relative intensity. Color coding: VA (red), VP (green), DA (blue), and DP (yellow) thalamus. Some beams were not drawn due to unrecorded head-transducer contact coordinates: DP of P29 and VP of P44. VA: ventral anterior thalamus; VP: ventral posterior thalamus; DA: dorsal anterior thalamus; DP: dorsal posterior thalamus.

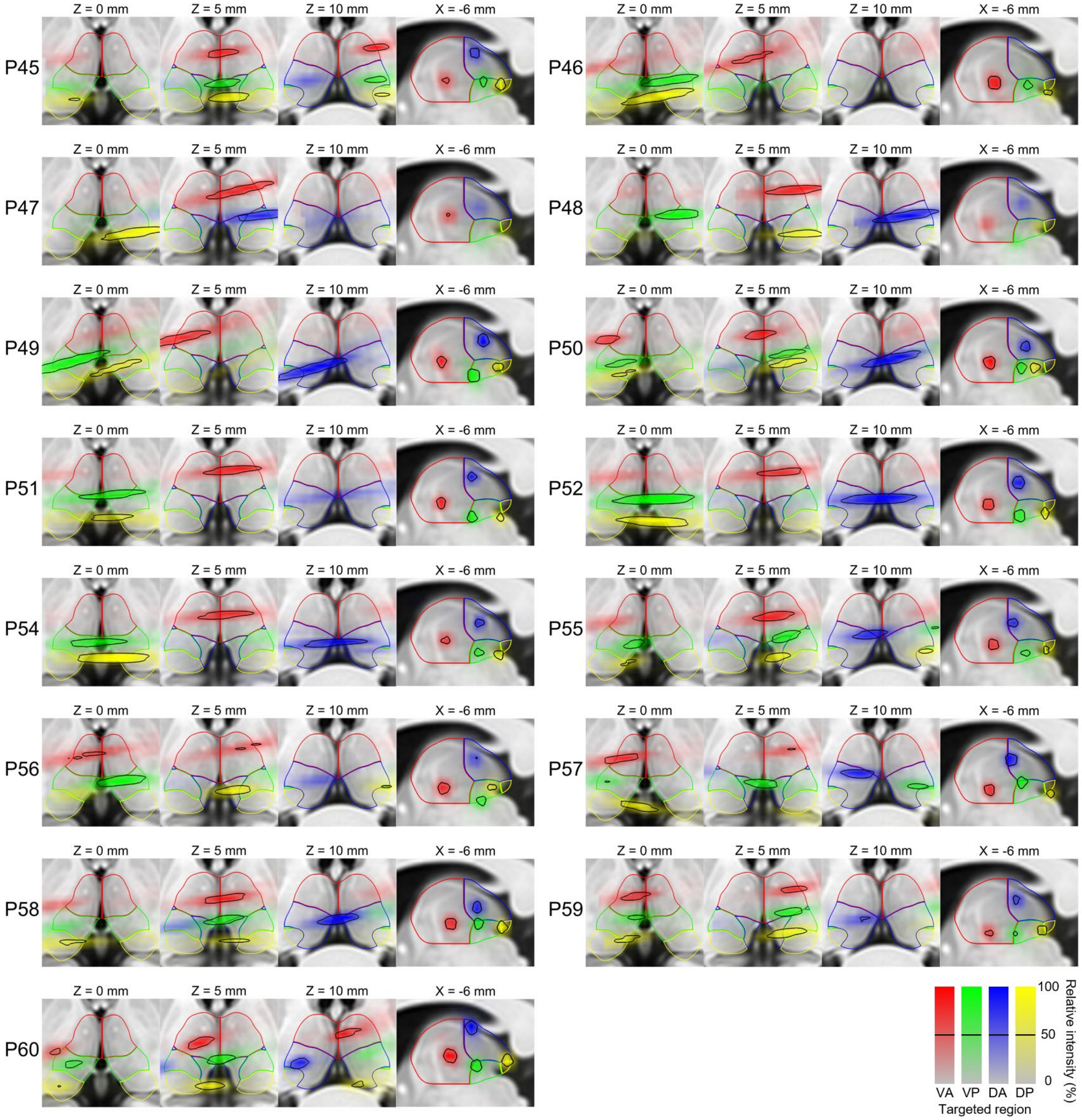


**Supplementary Fig. 10: Individual-level mosaic views of beam trajectories for participants P45 to P60.** Black contours indicate 50% (-3 dB) relative intensity. Color coding: VA (red), VP (green), DA (blue), and DP (yellow) thalamus. Some beams were not drawn due to reasons as follows: unrecorded head-transducer contact coordinates: DA of P46; beam simulated but not visible at our chosen cross-sections: VP of P47. VA: ventral anterior thalamus; VP: ventral posterior thalamus; DA: dorsal anterior thalamus; DP: dorsal posterior thalamus.

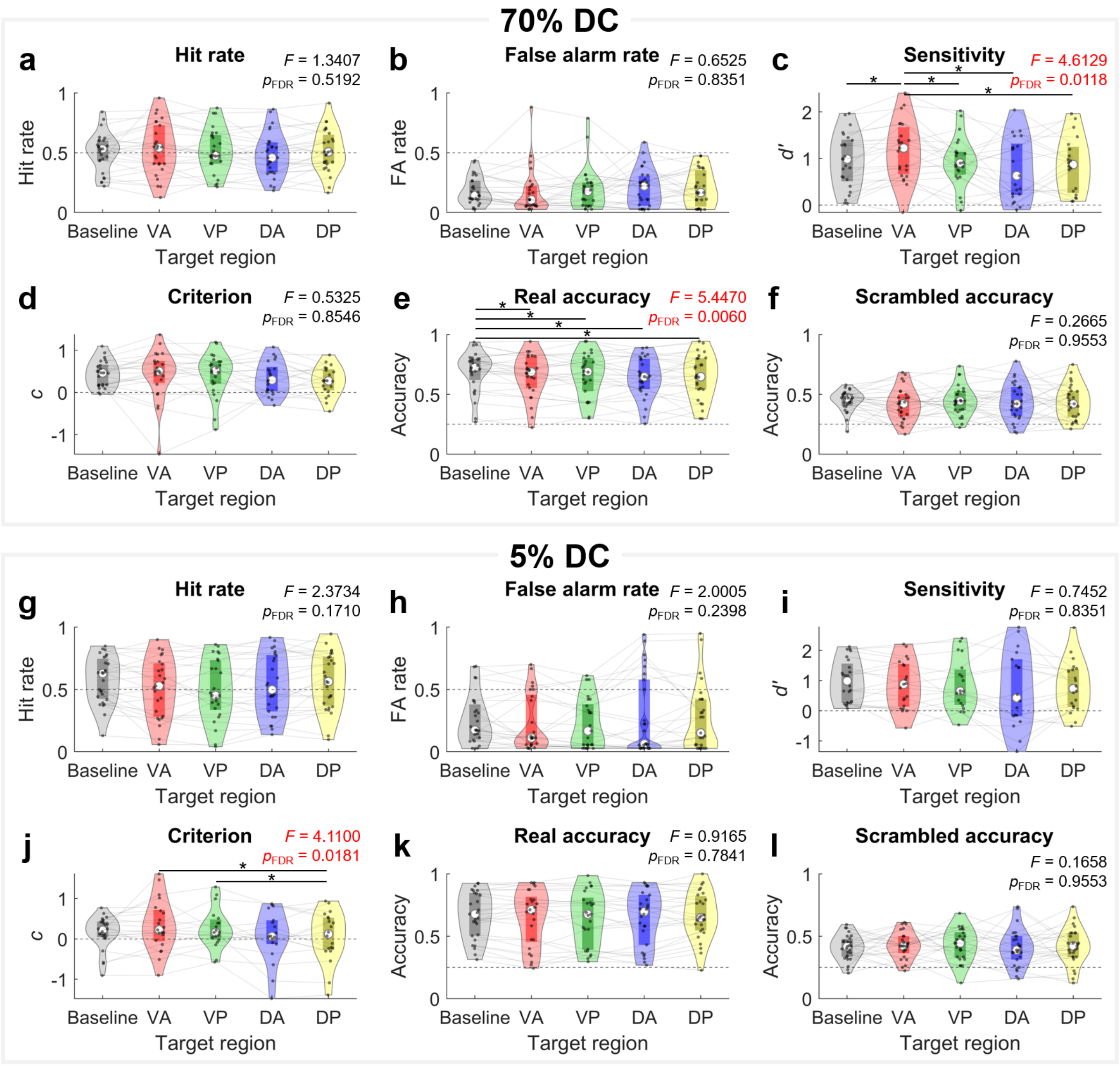


**Supplementary Fig. 11: Entire target-specific perceptual outcomes during human thalamic LIFU administration.** Perceptual outcomes measured at **a-f** 70% and **g-l** 5% duty cycle, including **a, g** hit rate, **b, h** false alarm rate, **c, i** sensitivity *d′*, **d, j** criterion *c*, **e, k** categorization accuracy for real images, and **f, l** categorization accuracy for scrambled images. Baseline values are outcomes averaged across two baselines. The *F*-values and FDR-corrected *p*-values of omnibus test are provided. Post-hoc pairwise comparisons were performed only for FDR-corrected $p < 0.05$. Asterisks denote significant pairwise differences (FDR-corrected $p < 0.05$). Boxes indicate interquartile ranges. Median values are marked by white circles. Dashed lines are reference values for each measure: hit rate of 0.5 (a, g), no discrimination of $d' = 0$ (c, i), no bias of $c = 0$ (d, j), and chance accuracy of 0.25 (e, f, k, l). Sample sizes are as follows: (a,b,e,f) Baseline: $n$ = 27; VA: $n$ = 26; VP: $n$ = 26; DA: $n$ = 27; DP: $n$ = 27; (c,d) Baseline: $n$ = 25; VA: $n$ = 23; VP: $n$ = 23; DA: $n$ = 21; DP: $n$ = 20; (g,h,k,l) Baseline: $n$ = 27; VA: $n$ = 25; VP: $n$ = 25; DA: $n$ = 26; DP: $n$ = 27; (i,j) Baseline: $n$ = 24; VA: $n$ = 21; VP: $n$ = 19; DA: $n$ = 18; DP: $n$ = 20. DC: duty cycle; FA: false alarm; VA: ventral anterior thalamus; VP: ventral posterior thalamus; DA: dorsal anterior thalamus; DP: dorsal posterior thalamus. Statistics and source data are provided as a Source Data file.

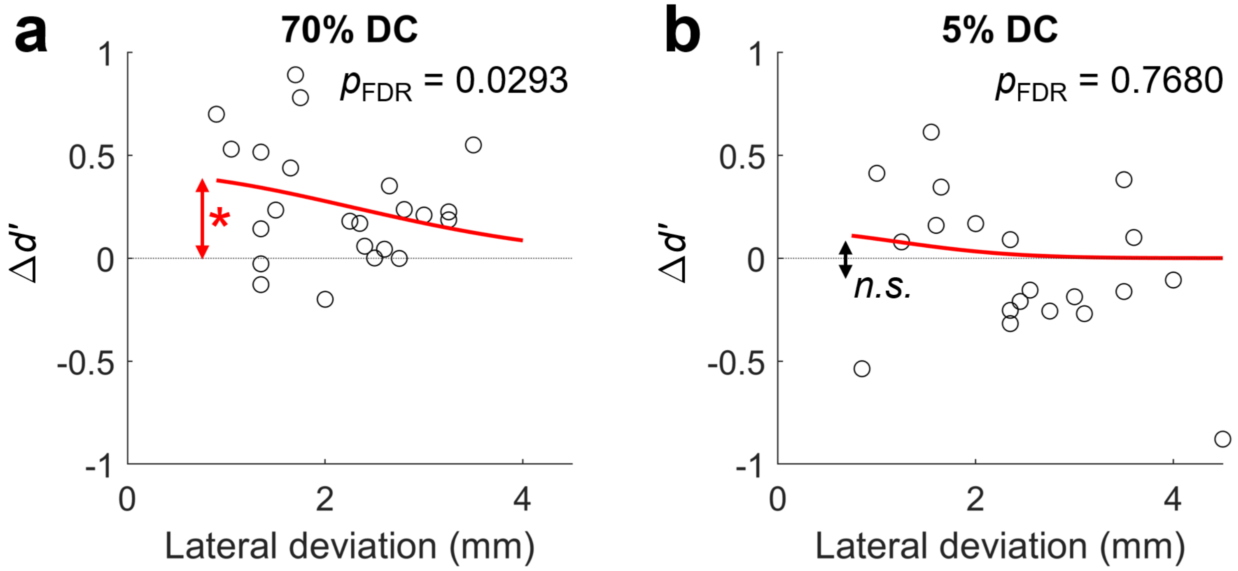


**Supplementary Fig. 12: Sensitivity change during VA thalamus sonication in relation to lateral deviation.** Baseline-subtracted sensitivity *d′* during VA thalamus sonication at **a** 70% ($n$ = 23) and **b** 5% duty cycles ($n$ = 20), plotted against lateral deviation. Red curves are best fit to a Gaussian curve ($y = c_1\, exp(-x^2/c_2)$). Asterisk indicates significant non-zero height (i.e., parameter $c_1$) of the fitted Gaussian curves (FDR-corrected *p*-values denoted at top-left). The *n.s.* denotes non-significance. Fitting statistics and source data are provided as a Source Data file.

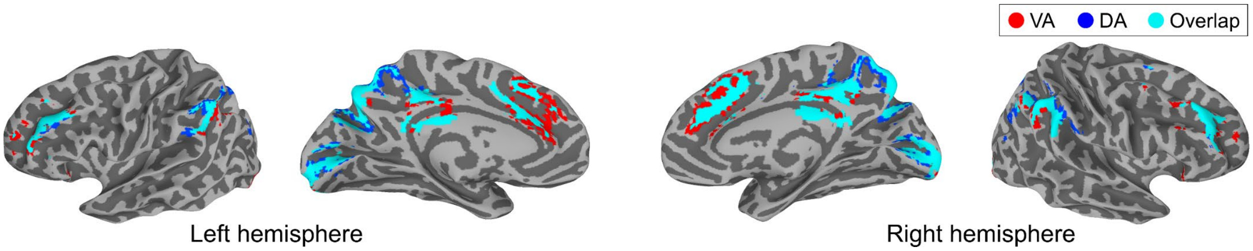


**Supplementary Fig. 13: Functional connectivity of VA (ventral anterior) and DA (dorsal anterior) thalamus with the cortex.** Cortical voxels with top 10% strongest connectivity with VA and DA thalamus. Overlapping voxels are highlighted as cyan.

**Supplementary Table 1: A report summary based on ITRUSST reporting guidelines.**

| **Transducer and drive system description** | | | | |
|---|---|---|---|---|
| Transducer manufacturer and model number | BrainSonix, BXPulsar 1002 | | | |
| Transducer center frequency | 659.6 kHz | | | |
| Transducer geometry | Aperture: 61.5 mm | | | |
| Drive system components | All-in-one system (POC-155, Advantech) | | | |
| **Drive system settings** | | | | |
| Operating frequency | 650 kHz | | | |
| Output level settings | 0.72 W $cm^{-2}$ ($I_{spta.3}$) | | | |
| Focal position settings | Fixed | | | |
| **Free field acoustic parameters** | | | | |
| Spatial-peak pressure amplitude | 197 kPa (70% DC) and 710 kPa (5% DC) | | | |
| Position of spatial-peak pressure amplitude | 80.7 mm | | | |
| Size of focal volume | 30.3 mm (length), 3.3 mm (width) | | | |
| Description | Focal depth was validated by Blatek Industries, Inc. with ASTM E1065/E1065M Standard guidelines. Focal volume size was measured with a bilaminar membrane hydrophone with a 0.4 mm diameter active receive aperture (data provided by BrainSonix Corporations). | | | |
| **Pulse timing parameters** | | | | |
| | Duration | Ramp duration | Ramp shape | Repetition interval/frequency |
| 70% DC Pulse<br>Pulse train<br>Pulse train repeat | 70 ms<br>30 s<br>11.5 min | 0<br>0<br>0 | Rectangular<br>Rectangular<br>Rectangular | 100 ms/10 Hz<br>60 s/0.017 Hz |
| 5% DC Pulse<br>Pulse train<br>Pulse train repeat | 5 ms<br>30 s<br>11.5 min | 0<br>0<br>0 | Rectangular<br>Rectangular<br>Rectangular | 100 ms/10 Hz<br>60 s/0.017 Hz |
| ***In situ* estimates of exposure parameters** | | | | |
| Estimated *in situ* pressure amplitude at the target | 72.6 ± 4.2 kPa (70% DC, mean ± SD)<br>258.2 ± 18.3 kPa (5% DC, mean ± SD) | | | |
| Estimated *in situ* mechanical index | 0.20 (70% DC); 0.77 (5% DC) | | | |
| Estimated temperature rise | 0.23 ± 0.04 °C (mean ± SD) | | | |
| Description | Pressure and temperature rise were simulated *in silico* on BabelBrain.<br>Mechanical index was calculated from $P_{r.3}$, without considering skull attenuation. | | | |

**Supplementary Table 2: Test data report of the transducer used in the study.**

| Transducer information | |
|---:|---|
| Part number | AT29823 |
| Description | BrainSonix, 650 kHz, 61.5 mm diameter, 80 mm focal depth, 10 ft cable (BNC) |
| **Test data** | |
| Serial number | 0919-5441 |
| Test time | 7:14 AM 12/29/2021 |
| Sensitivity | 588 mV |
| Frequency bandwidth | 28% |
| -20dB pulse length | 9.201 µs |
| Center frequency | 659.6 kHz |
| Test target | 1” steel @ focal |
| Test path length | 80.7 mm |
| **Equipment used** | |
| Oscilloscope model | LeCroy WS452 |
| Oscilloscope calibration due date | 6/30/2022 |
| Pulser model | 5072PR |
| Pulser calibration due date | 07/19/2022 |
| **Pulser/receiver settings** | |
| Energy level | 4 |
| Damping | 3 (50 Ohm) |
| Gain | -23 dB |

| Signal waveform | Frequency spectrum |
|---|---|
| 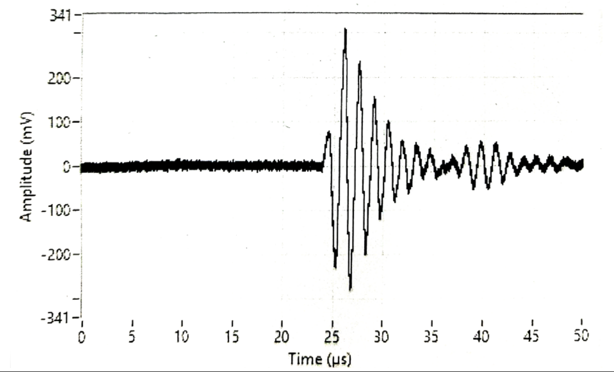  | 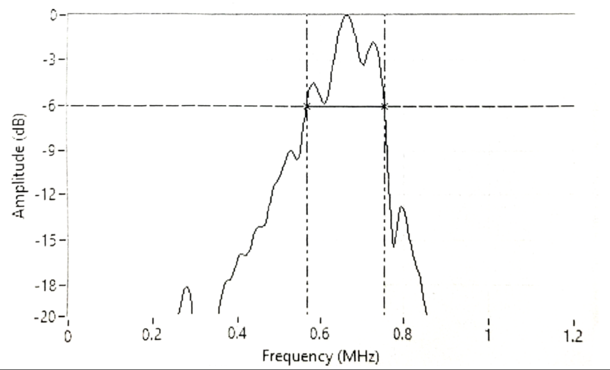  |

**Supplementary Table 3: Omnibus ANOVA test replicated with individual baselines without averaging.**

| Baseline-1 | | | | | | |
|---|---|---|---|---|---|---|
| **DC** | **Outcome** | **$df_1$** | **$df_2$** | ***F*-value** | ***p*-value** | **FDR-corrected *p*-value** |
| **70%** | Hit rate | 4 | 108 | 1.1169 | 0.3524 | 0.7049 |
| | FA rate | 4 | 109 | 0.7117 | 0.5856 | 0.8378 |
| | Sensitivity *d′* | 4 | 90 | 4.2123 | 0.0036 | 0.0215 |
| | Criterion *c* | 4 | 89 | 0.4816 | 0.7492 | 0.8990 |
| | Real image accuracy | 4 | 107 | 4.2912 | 0.0029 | 0.0215 |
| | Scrambled image accuracy | 4 | 111 | 0.1927 | 0.9418 | 0.9657 |
| **5%** | Hit rate | 4 | 104 | 1.6848 | 0.1591 | 0.3970 |
| | FA rate | 4 | 104 | 1.6581 | 0.1654 | 0.3970 |
| | Sensitivity *d′* | 4 | 78 | 0.6504 | 0.6283 | 0.8378 |
| | Criterion *c* | 4 | 77 | 3.0564 | 0.0215 | 0.0861 |
| | Real image accuracy | 4 | 103 | 0.7181 | 0.5814 | 0.8378 |
| | Scrambled image accuracy | 4 | 105 | 0.1431 | 0.9657 | 0.9657 |
| **Baseline-2** | | | | | | |
| **DC** | **Outcome** | **$df_1$** | **$df_2$** | ***F*-value** | ***p*-value** | **FDR-corrected *p*-value** |
| **70%** | Hit rate | 4 | 105 | 1.2346 | 0.3007 | 0.6015 |
| | FA rate | 4 | 107 | 0.6043 | 0.6604 | 0.8075 |
| | Sensitivity *d′* | 4 | 84 | 3.4286 | 0.0120 | 0.0480 |
| | Criterion *c* | 4 | 84 | 0.4939 | 0.7402 | 0.8075 |
| | Real image accuracy | 4 | 104 | 3.8119 | 0.0062 | 0.0480 |
| | Scrambled image accuracy | 4 | 108 | 0.4955 | 0.7390 | 0.8075 |
| **5%** | Hit rate | 4 | 103 | 2.1152 | 0.0842 | 0.2527 |
| | FA rate | 4 | 103 | 1.8738 | 0.1207 | 0.2896 |
| | Sensitivity *d′* | 4 | 76 | 0.7877 | 0.5367 | 0.8075 |
| | Criterion *c* | 4 | 75 | 3.6633 | 0.0088 | 0.0480 |
| | Real image accuracy | 4 | 102 | 0.7263 | 0.5760 | 0.8075 |
| | Scrambled image accuracy | 4 | 104 | 0.1733 | 0.9516 | 0.9516 |

Significances (FDR-corrected $p < 0.05$) are highlighted in red.

**Supplementary Table 4: FDR-corrected *p*-values for the effects of six confound variables in omnibus ANOVA.**

| DC | Outcome | Age | Sex | Transducer noise awareness | Lateral deviation | Attenuation ratio | Sonication order |
|---|---|---|---|---|---|---|---|
| 70% | Hit rate | 0.9986 | 0.9934 | 0.7846 | 0.2514 | 0.9285 | 0.9180 |
| | FA rate | 0.9986 | 0.9934 | 0.7666 | 0.9910 | 0.9285 | 0.7460 |
| | Sensitivity *d′* | 0.9986 | 0.9934 | 0.7666 | 0.0898 | 0.9285 | 0.7460 |
| | Criterion *c* | 0.9986 | 0.9934 | 0.7666 | 0.8964 | 0.9285 | 0.7460 |
| | Real image accuracy | 0.9986 | 0.9934 | 0.7666 | 0.4454 | 0.1975 | 0.7460 |
| | Scrambled image accuracy | 0.9986 | 0.9934 | 0.7666 | 0.9910 | 0.9285 | 0.9249 |
| 5% | Hit rate | 0.9986 | 0.9934 | 0.7846 | 0.9910 | 0.9285 | 0.7460 |
| | FA rate | 0.9986 | 0.9934 | 0.7666 | 0.0898 | 0.9285 | 0.7460 |
| | Sensitivity *d′* | 0.9986 | 0.9934 | 0.7666 | 0.0719 | 0.9285 | 0.7460 |
| | Criterion *c* | 0.9986 | 0.9934 | 0.7666 | 0.0687 | 0.9285 | 0.7460 |
| | Real image accuracy | 0.9986 | 0.9934 | 0.7666 | 0.9910 | 0.9285 | 0.7460 |
| | Scrambled image accuracy | 0.9986 | 0.9934 | 0.7666 | 0.9910 | 0.9285 | 0.9180 |

Full statistics including *F*-values and uncorrected *p*-values are provided in Source Data.

**Supplementary Table 5: Post-hoc pairwise comparison results.**

| Sensitivity *d'* at 70% DC | | | | |
|---|---|---|---|---|
| **Comparison** | ***F*-value** | ***p*-value** | **FDR-corrected *p*-value** | **Direction** |
| Baseline vs. VA | 7.3460 | 0.0079 | 0.0246 | VA > Baseline |
| Baseline vs. VP | 0.4758 | 0.4919 | 0.7379 | - |
| Baseline vs. DA | 0.0003 | 0.9866 | 0.9866 | - |
| Baseline vs. DP | 0.0467 | 0.8293 | 0.9135 | - |
| VA vs. VP | 7.2724 | 0.0082 | 0.0246 | VA > VP |
| VA vs. DA | 13.7511 | 0.0003 | 0.0034 | VA > DA |
| VA vs. DP | 9.9654 | 0.0021 | 0.0119 | VA > DP |
| VP vs. DA | 0.8079 | 0.3709 | 0.6181 | - |
| VP vs. DP | 0.3813 | 0.5383 | 0.7530 | - |
| DA vs. DP | 0.0875 | 0.7680 | 0.8861 | - |
| **Real image categorization accuracy at 70% DC** | | | | |
| **Comparison** | ***F*-value** | ***p*-value** | **FDR-corrected *p*-value** | **Direction** |
| Baseline vs. VA | 9.6224 | 0.0024 | 0.0119 | Baseline > VA |
| Baseline vs. VP | 12.2472 | 0.0007 | 0.0049 | Baseline > VP |
| Baseline vs. DA | 16.6370 | 0.0001 | 0.0012 | Baseline > DA |
| Baseline vs. DP | 19.3209 | 0.0000 | 0.0007 | Baseline > DP |
| VA vs. VP | 0.2969 | 0.5869 | 0.7655 | - |
| VA vs. DA | 2.0056 | 0.1593 | 0.2986 | - |
| VA vs. DP | 2.7777 | 0.0982 | 0.1963 | - |
| VP vs. DA | 0.6387 | 0.4257 | 0.6722 | - |
| VP vs. DP | 1.4150 | 0.2365 | 0.4174 | - |
| DA vs. DP | 0.1061 | 0.7452 | 0.8861 | - |
| **Criterion *c* at 5% DC** | | | | |
| **Comparison** | ***F*-value** | ***p*-value** | **FDR-corrected *p*-value** | **Direction** |
| Baseline vs. VA | 4.6718 | 0.0333 | 0.0832 | VA > Baseline |
| Baseline vs. VP | 4.1937 | 0.0435 | 0.0931 | VP > Baseline |
| Baseline vs. DA | 0.0347 | 0.8526 | 0.9135 | - |
| Baseline vs. DP | 0.1309 | 0.7183 | 0.8861 | - |
| VA vs. VP | 0.0012 | 0.9725 | 0.9866 | - |
| VA vs. DA | 5.6630 | 0.0194 | 0.0529 | VA > DA |
| VA vs. DP | 8.3334 | 0.0049 | 0.0182 | VA > DP |
| VP vs. DA | 4.3694 | 0.0394 | 0.0909 | VP > DA |
| VP vs. DP | 9.3500 | 0.0029 | 0.0125 | VP > DP |
| DA vs. DP | 0.3560 | 0.5522 | 0.7530 | - |

Directions of change are denoted for uncorrected $p < 0.05$. Red indicates FDR-corrected $p < 0.05$. DC: duty cycle; VA: ventral anterior thalamus; VP: ventral posterior thalamus; DA: dorsal anterior thalamus; DP: dorsal posterior thalamus.

**Supplementary Information References**

1. Podvalny, E., Flounders, M. W., King, L. E., Holroyd, T. & He, B. J. A dual role of prestimulus spontaneous neural activity in visual object recognition. *Nat Commun* **10**, 3910 (2019).

2. Wu, Y., Podvalny, E., Levinson, M. & He, B. J. Network mechanisms of ongoing brain activity's influence on conscious visual perception. *Nat Commun* **15**, 5720 (2024).
